# Deciphering the Influence of Socioeconomic Status on Brain Structure: Insights from Mendelian Randomization

**DOI:** 10.1101/2024.03.13.584410

**Authors:** Charley Xia, Yuechen Lu, Zhuzhuoyu Zhou, Mattia Marchi, Hyeokmoon Kweon, Yuchen Ning, David C. M. Liewald, Emma L. Anderson, Philipp D. Koellinger, Simon R. Cox, Marco P. Boks, W. David Hill

**Affiliations:** Lothian Birth Cohort studies, University of Edinburgh, 7 George Square, Edinburgh EH8 9JZ, UK; Department of Psychology, School of Philosophy, Psychology and Language Sciences, University of Edinburgh, 7 George Square, Edinburgh EH8 9JZ, UK; Department of Biomedical, Metabolic and Neural Sciences, University of Modena and Reggio Emilia, Via Giuseppe Campi, 287 – 41125 Modena, Italy; Department of Mental Health and Addiction Services, Azienda USL-IRCCS di Reggio Emilia, Reggio Emilia, Italy; Department of Psychiatry, Brain Center University Medical Center Utrecht, University Utrecht, Utrecht, The Netherlands; Department of Economics, School of Business and Economics, Vrije Universiteit Amsterdam, De Boelelaan 1105, 1081 HV Amsterdam, The Netherlands; MRC Integrative Epidemiology Unit, Bristol Medical School, University of Bristol, Oakfield House, Oakfield Grove, Bristol, BS8 2BN, United Kingdom; Division of Psychiatry, Faculty of Brain Sciences, University College London, London, United Kingdom

## Abstract

Socioeconomic status (SES) influences physical and mental health, however its relation with brain structure is less well documented. Here, we examine the role of SES on brain structure using Mendelian randomisation. First, we conduct a multivariate genome-wide association study of SES using individual, household, and area-based measures of SES, with an effective sample size of n=893,604. We identify 469 loci associated with SES and distil these loci into those that are common across measures of SES and those specific to each indicator. Second, using an independent sample of ∼35,000 we provide evidence to suggest that total brain volume is a causal factor in higher SES, and that SES is protective against white matter hyperintensities as a proportion of intracranial volume (WMHicv). Third, we find evidence that whilst differences in cognitive ability explain some of the causal effect of SES on WMHicv, differences in SES still afford a protective effect against WMHicv, independent of that made by cognitive ability.

## Introduction

Socioeconomic status (SES) is a multi-dimensional construct influencing, and influenced by, multiple physical, socio-cultural, and environmental factors. Differences in SES are a determining factor of health where those from more advantaged backgrounds have a higher level of physical health, mental health and psychiatric conditions, are less likely to receive a dementia diagnosis, and live longer lives^1–4^. These inequalities in physical health, and mental health are present across different indicators of SES and have been found for occupation, income, educational attainment, and measures of social deprivation^1,5–7^. The communality of such findings highlights the need to examine the influence of SES using a multifactorial approach to examine the causes and consequences of differences in SES.

As with any other quantitative trait, such as height or weight, differences in SES have a heritable component^8^ meaning that genetic differences within a population will covary with phenotypic differences. However, and unlike traits such as height or weight, such genetic differences associated with SES are unlikely to form part of a biological pathway from gene to phenotype directly, but are more likely the result of a phenotypic pathway, known as vertical pleiotropy^9^, where a number of traits (which are themselves heritable) contribute towards differences in SES^10–12^. As such, the heritability of SES is not static, and differences between societies can result in differences in the heritable traits that give rise to the observed differences in SES^13,14^.

Despite the genetically heterogeneous nature of SES, genome-wide association studies (GWAS) examining indicators of SES such as measures of income^11^, educational attainment^15^, and social deprivation^8^ have identified hundreds of associated genetic loci. These genetic indicators of SES are also linked to physical health outcomes, indicative of a common genetic aetiology between SES and physical health^8,11,15^. Furthermore, psychiatric traits including schizophrenia, major depressive disorder, and attention deficit hyperactivity disorder, as well as neurological disorders such as Alzheimer’s disease, early-onset stroke, and intra-cerebral haemorrhage also share genetic effects with measures of SES^16^. As with genetic influences that act on SES, these links between SES and brain-related health outcomes may themselves include other phenotypes such as neuroanatomy^11^.

Genetic links between SES and brain morphology have been identified previously where genes highly expressed in the brain and both neuronal and glial cells are enriched for their associations with both income^11^ and educational attainment^17^. Furthermore, strong genetic correlations are found between indicators of SES and brain morphology where a genetic correlation of *r_g_* = 0.34, SE = 0.07, P =1.2×10^-6^ has been identified between intracranial volume and educational attainment^18^, and loci associated with cognitive ability^19^ are found to be overrepresented in the associated loci from a GWAS of income^11^. These genetic links between SES and brain-based phenotypes have been explored using Mendelian randomisation (MR) to examine the direction of causality between them. For example, evidence of bidirectional causal effects was found between poverty (n = 668,288) and mental illness^20^ using MR.

However, the following are some fundamental gaps in our understanding of the relationship between SES and brain structure. First, do different indicators of SES confer the different levels of risk or is SES best captured using a single factor? Second, is there evidence for causality in the relationship between SES and brain morphology? Third, to what extend does differences in cognitive ability explain the relationship between SES and brain morphology?

Importantly, the use of brain morphology as an outcome in MR can allow for the risk factors of late-life cognitive function that act on cognitive decline in adulthood to be distinguished from those that differentiate the trajectory of cognitive growth through childhood. The importance of which is underscored in the context of dementia which, whilst typically diagnosed using cognitive tests such as the Mini-Mental State Examination^21^, is distinguished from other neurodevelopmental disorders (such as intellectual disability) by a progressive later-life loss of cognitive ability that affects daily life^22^. As such, risk of dementia can be seen to be composed of two components: cognitive development influencing the level of cognitive function prior to the onset of cognitive decline and, the rate at which decline occurs. Whilst large GWAS of cognitive decline are currently lacking, MR combined with GWAS conducted on frank indictors of brain ageing, such as white matter hyperintensities^23^, can be used to identify potentially modifiable risk factors causal in brain ageing.

In the current study, we combine multivariate analysis with MR to examine the bidirectional effects between SES and brain morphology and to examine likely heritable traits that are captured by measures of SES, and to identify potentially modifiable risk factors of age-related brain change associated with cognitive development and cognitive decline. First, we perform a common-factor model multi-variate GWAS of four indicators of SES: occupational prestige (OP, n = 279,644), household income (HI, n = 488,233), educational attainment (EA, n = 753,152), and social deprivation (SD, n = 440,350) for an effective size of 893,604 participants. The use of these four measures in a multivariate framework allows for the assessment of heterogeneous effects across indicators of SES in conjunction with an investigation of common genetic effects that act on the individual, as well as the household, and geographical area in which one resides. Thus, genetic effects can be categorised as common across measures of SES or unique to specific indicators. Second, to examine the bidirectional causal effects of SES on brain structure we use two-sample MR on 13 brain imaging phenotypes sourced from an independent sample of ∼36,000 UKB participants. Finally, we examine the role of cognitive ability on the links between SES and brain morphology as one of the heritable traits that is captured by GWAS conducted on the general factor of SES and these four indicators.

## Results

### Study design and implementation

Genome-wide association study (GWAS) data sets were used to identify instrumental variables for five exposures. These were four measures of SES (occupational prestige, household income, educational attainment, and social deprivation), and one cognitive exposure (cognitive ability). GWASs were also performed in an independent sample on thirteen MRI phenotypes (total brain volume, TBV; grey matter volume, GM; normal appearing white matter, NAWM; white matter hyperintensity volume, WMH; TBV as a proportion of intracranial volume, TBVicv; GM as a proportion of intracranial volume, GMicv; white matter volume as a proportion of intracranial volume, WMicv; WMH as a proportion of intracranial volume, WMHicv; a general factor of brain white matter tract fractional anisotropy, gFA; a general factor of brain white matter tract mean diffusivity, gMD; a general factor of brain white matter tract intracellular volume fraction, gIVCF; a general factor of brain white matter tract isotropic volume fraction, gISOVF; a general factor of brain white matter tract orientation dispersion, gOD) capturing different aspects of brain morphology.

### Phenotypic and genetic structure of the four indices of socioeconomic status

The phenotypic correlations between the four indices of SES (**Supplementary Table 1**) were all significant and ranged from *r* = 0.062 – 0.484 (mean = 0.268, SE range = 0.00143 – 0.00185, P < 10^-322^). A confirmatory factor model with a single common factor fit the phenotypic data poorly (χ^2^(2) = 8530.202, *p* < 0.001; SRMR=0.047; CFI=0.932; RMSEA = 0.131, TLI=0.795, **Figure 1** & **Table 1**). The common factor explained 31.373% of the phenotypic variance across each of the four indices of SES.

**Figure 1.**
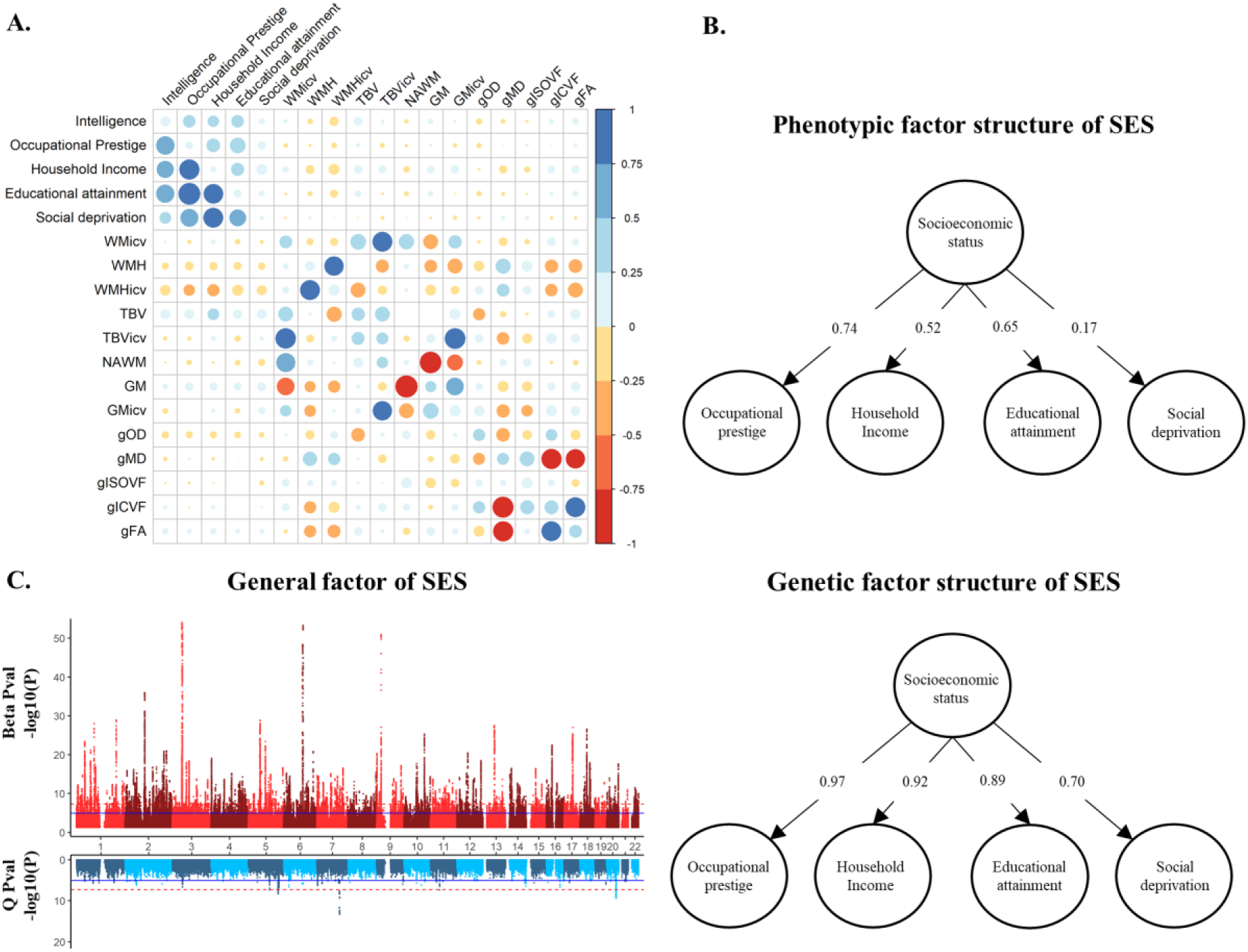
**A.** Showing the phenotypic and genetic correlations between the variables used. The lower diagonal shows the genetic correlations whereas the upper diagonal shows the phenotypic correlations. The diagonal shows the heritability estimates. Colour and size are used to illustrate the magnitude and directions of the correlations. Both heritability and genetic correlations were derived using LDSC implemented in Genomic SEM. Tabulated values are shown in **Supplementary Tables 1-3.** Social deprivation scores were reversed to facilitate a comparison with the other measures of SES. **B.** Showing the standardised phenotypic (upper) and genetic (lower) factor solutions for the covariance structure across the four indices of SES examined in the total sample. Social deprivation scores were again reversed. **C.** A miami plot of the general factor of SES. The X axis indicates chromosome and the y axis shows the –log(10) p value of each SNP with the upper section describing its association with the general factor of SES where the lower shows the p value for the heterogeneity Q statistics. TBV, total brain volume; GM, grey matter volume; WMH, white matter hyperintensity volume; TBVicv, TBV as a proportion of intracranial volume; GMicv, GM as a proportion of intracranial volume; WMicv, white matter volume as a proportion of intracranial volume; WMHicv, WMH as a proportion of intracranial volume; gFA, a general factor of brain white matter tract fractional anisotropy; gMD, a general factor of brain white matter tract mean diffusivity; gIVCF, a general factor of brain white matter tract intracellular volume fraction; gISOVF, a general factor of brain white matter tract isotropic volume fraction; NAWM, normal appearing white matter; gOD, a general factor of brain white matter tract orientation dispersion.

**Table 1.**
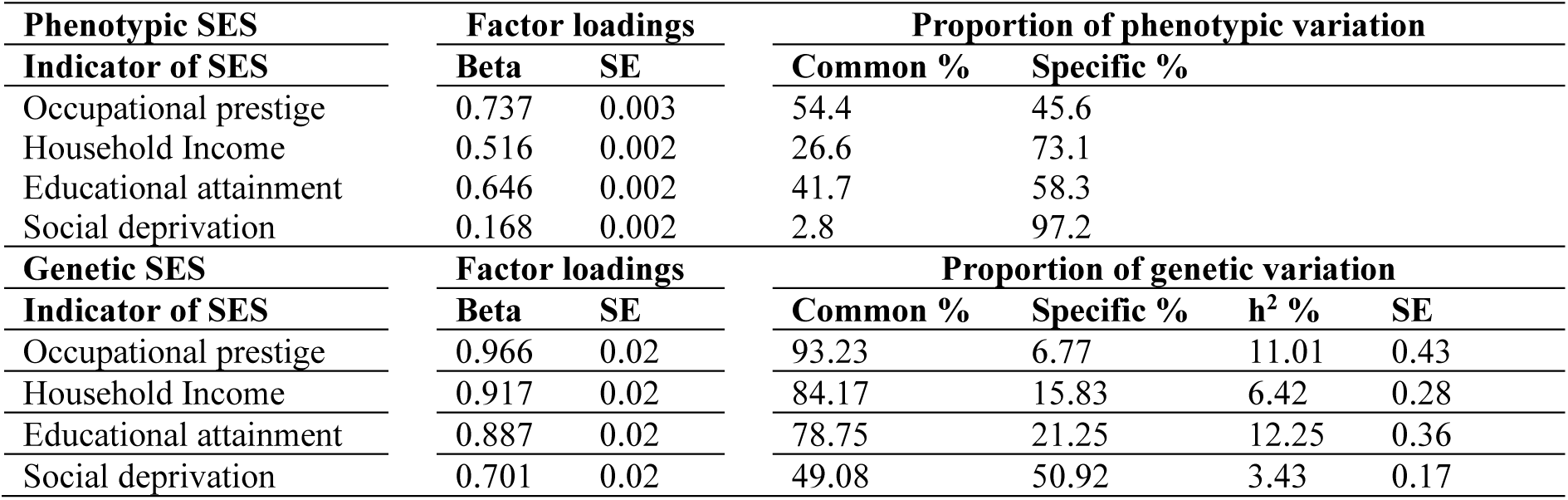
Showing the standardised factor loadings for each of the four indicators of SES in the total sample. The direction of social deprivation was reversed so that all scores indicate a greater level of SES across the four indicators used. The upper portion shows the phenotypic structure of SES where the bottom portion shows the genetic structure of SES. Common and specific, by definition sum to 100%, but for the genetic structure this indicates the proportion from common and specific sources that contribute to the total heritability. The total heritability was derived using LDSC implemented in genomic SEM.

Using LDSC^24^ on each of the GWASs conducted on the indices of SES (occupational prestige, household income, educational attainment, and social deprivation), a significant heritable component was captured explaining between 3.5% - 13% of trait variation (**Supplementary Table 2**). LDSC intercepts were consistently close to 1 for each SES measure indicating that polygenicity, rather than population stratification or other factors, explained the inflation in GWAS association test statistics (**Supplementary Table 2**).

Strong genetic correlations between indicators of SES (mean *r_g_* = 0.761, range *r_g_* = 0.563 – 0.963, SE range = 0.011 - 0.026) were observed (**Supplementary Table 3**). The moderate phenotypic correlations but large genetic correlations indicate that whilst each measure of SES captures a different environmental component, they each draw upon similar heritable traits. This was confirmed by extracting a general genetic factor of SES using genomic structural equation modelling (Genomic SEM^25^, **Figure 1 B** & **Table 1**) where, in contrast to the phenotypic data, a single factor explained the covariance across the genetic data sets well (χ2(2) = 184.737, *p* = 7.67×10^−41^; SRMR=0.051; CFI=0.988). The general genetic factor of SES captured on average 76.30% of the genetic variance in each indicator of SES with the proportion being consistent across occupational prestige, household income, and educational attainment (>75%), with the lowest being social deprivation where the general factor captured 49.08% (**Supplementary Table 4**). This general factor of SES was then regressed onto 7,462,726 SNPs to derive genome wide associations with SES. Furthermore, to differentiate between SNPs whose loci are relevant to a general factor of SES from those whose patterns of association are inconsistent with a general factor, we derive genome-wide heterogeneity statistics using Genomic SEM^25^. The general genetic factor of SES had a h^2^ = 9.40% (SE = 0.20%), and showed little evidence of inflation in test statistics due to population stratification (LDSC intercept = 1.06, SE = 0.02). In order to attain independent groups to perform Two-sample MR a general factor of SES was also derived by omitting participants and their relatives who contributed MRI data. This resulted in an effective sample size of 665,662 participants. This general factor had a highly similar factor structure as the full data set (**Supplementary Table 4**) and a similar heritability (h^2^ = 11.3%, SE = 0.30%).

### Estimating bi-directional causal effects of SES on brain structure

Using Steiger filtering^26^ followed by two-sample Mendelian randomisation (MR)^27^ a higher SES was found to be a protective factor against WMHicv (β = −0.218, SE = 0.056, P = 8.63×10^−5^, **Table 2**, **Supplementary Table 5, Supplementary Figures 1 & 2**). The use of both MR-Egger and MR-PRESSO did not identify any horizontal pleiotropy and no significant heterogeneity was found (**Supplementary Table 5** & **Supplementary Table 6**). There was very little evidence of any other causal effects.

**Table 2.**
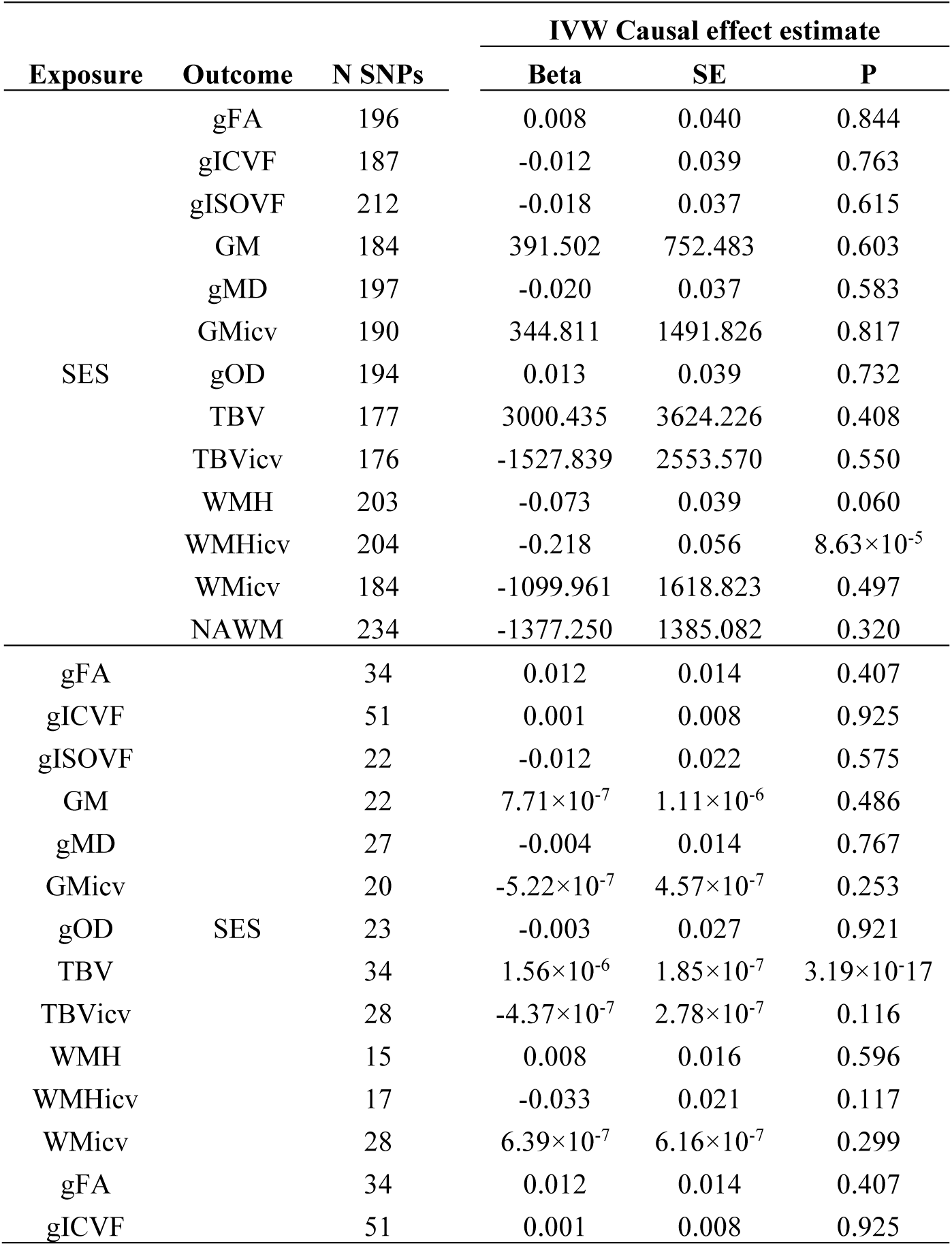
Showing the inverse variance weighted bi-directional causal effect estimate of socioeconomic status on brain structure. Abbreviations: TBV, total brain volume; GM, grey matter volume; WMH, white matter hyperintensity volume; TBVicv, TBV as a proportion of intracranial volume; GMicv, GM as a proportion of intracranial volume; WMicv, white matter volume as a proportion of intracranial volume; WMHicv, WMH as a proportion of intracranial volume; gFA, a general factor of brain white matter tract fractional anisotropy; gMD, a general factor of brain white matter tract mean diffusivity; gIVCF, a general factor of brain white matter tract intracellular volume fraction; gISOVF, a general factor of brain white matter tract isotropic volume fraction; gOD, a general factor of brain white matter tract orientation dispersion; SES, socio-economic status; SE, standard error; P, p-value; SNP: single nucleotide polymorphism.

In the reverse direction a greater total brain volume was associated with higher SES (β = 1.56×10^−6^, SE = 1.85×10^−7^, P = 3.19×10^−16^, **Table 2**, **Supplementary Table 5 & Supplementary Figure 3 & 4**). No horizontal pleiotropy was detected using MR-Egger or MR-PRESSO but significant heterogeneity was found (**Supplementary Table 5** & **Supplementary Table 6)**. None of the other structural brain measures showed evidence of being a causal factor in differences in SES.

### Estimating causal effects of specific indices of SES on brain structure

Following Steiger filtering, WMHicv’s were found to be a consequence of differences in household income, occupational prestige, and educational attainment. The direction of effect was the same across these indicators where a lower SES was found to be a causal factor in the increase of WMHicv.

Specifically, increases in household income and educational attainment had protective effects on WMHicv of β = −0.376, SE = 0.111, P = 0.001 and β = −0.593, SE = 0.128, P = 3.77×10^−6^ respectively (**Table 3, Supplementary Table 7 & Supplementary Figures 6-8**). There was no evidence that the results were influenced by horizontal pleiotropy using either MR-Egger^28^ or MR-PRESSO^29^ (**Supplementary Table 7**, **Supplementary Table 8**), and no evidence that the instruments were capturing heterogeneous effects.

**Table 3.**
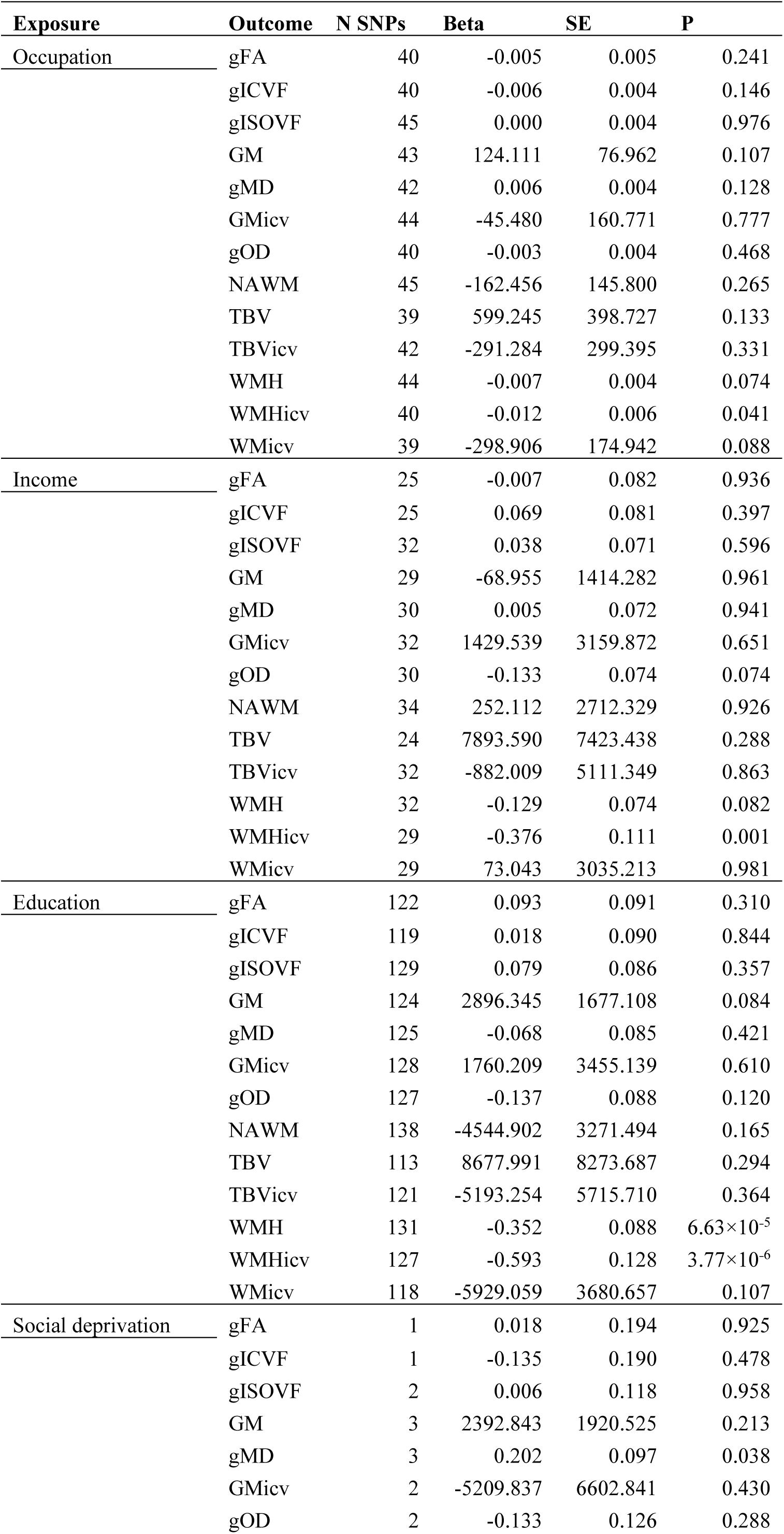
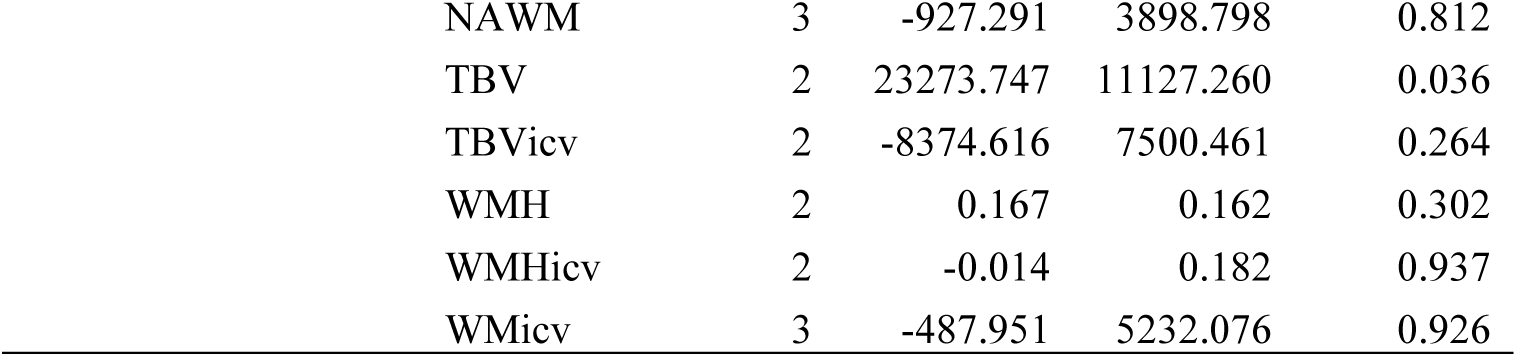
Showing the IVW causal effect estimate of each indicator of SES on brain measures. Beta weights are unstandardized and reflect the original unit of measure. Abbreviations: TBV, total brain volume; GM, grey matter volume; WMH, white matter hyperintensity volume; TBVicv, TBV as a proportion of intracranial volume; GMicv, GM as a proportion of intracranial volume; WMicv, white matter volume as a proportion of intracranial volume; WMHicv, WMH as a proportion of intracranial volume; gFA, a general factor of brain white matter tract fractional anisotropy; gMD, a general factor of brain white matter tract mean diffusivity; gIVCF, a general factor of brain white matter tract intracellular volume fraction; gISOVF, a general factor of brain white matter tract isotropic volume fraction; gOD,a general factor of brain white matter tract orientation dispersion. Note in the event that only one SNP was available a Wald ratio was used as an inverse-variance weighted model could not be derived.

| **Exposure** | **Outcome** | **N SNPs** | **Beta** | **SE** | **P** |
| --- | --- | --- | --- | --- | --- |
| <u>Occupation</u> | gFA | 40 | -0.005 | 0.005 | 0.241 |
|  | gICVF | 40 | -0.006 | 0.004 | 0.146 |
|  | gISOVF | 45 | 0.000 | 0.004 | 0.976 |
|  | GM | 43 | 124.111 | 76.962 | 0.107 |
|  | gMD | 42 | 0.006 | 0.004 | 0.128 |
|  | GMicv | 44 | -45.480 | 160.771 | 0.777 |
|  | gOD | 40 | -0.003 | 0.004 | 0.468 |
|  | NAWM | 45 | -162.456 | 145.800 | 0.265 |
|  | TBV | 39 | 599.245 | 398.727 | 0.133 |
|  | TBVicv | 42 | -291.284 | 299.395 | 0.331 |
|  | WMH | 44 | -0.007 | 0.004 | 0.074 |
|  | WMHicv | 40 | -0.012 | 0.006 | 0.041 |
|  | WMicv | 39 | -298.906 | 174.942 | 0.088 |
| <u>Income</u> | gFA | 25 | -0.007 | 0.082 | 0.936 |
|  | gICVF | 25 | 0.069 | 0.081 | 0.397 |
|  | gISOVF | 32 | 0.038 | 0.071 | 0.596 |
|  | GM | 29 | -68.955 | 1414.282 | 0.961 |
|  | gMD | 30 | 0.005 | 0.072 | 0.941 |
|  | GMicv | 32 | 1429.539 | 3159.872 | 0.651 |
|  | gOD | 30 | -0.133 | 0.074 | 0.074 |
|  | NAWM | 34 | 252.112 | 2712.329 | 0.926 |
|  | TBV | 24 | 7893.590 | 7423.438 | 0.288 |
|  | TBVicv | 32 | -882.009 | 5111.349 | 0.863 |
|  | WMH | 32 | -0.129 | 0.074 | 0.082 |
|  | WMHicv | 29 | -0.376 | 0.111 | 0.001 |
|  | WMicv | 29 | 73.043 | 3035.213 | 0.981 |
| <u>Education</u> | gFA | 122 | 0.093 | 0.091 | 0.310 |
|  | gICVF | 119 | 0.018 | 0.090 | 0.844 |
|  | gISOVF | 129 | 0.079 | 0.086 | 0.357 |
|  | GM | 124 | 2896.345 | 1677.108 | 0.084 |
|  | gMD | 125 | -0.068 | 0.085 | 0.421 |
|  | GMicv | 128 | 1760.209 | 3455.139 | 0.610 |
|  | gOD | 127 | -0.137 | 0.088 | 0.120 |
|  | NAWM | 138 | -4544.902 | 3271.494 | 0.165 |
|  | TBV | 113 | 8677.991 | 8273.687 | 0.294 |
|  | TBVicv | 121 | -5193.254 | 5715.710 | 0.364 |
| | WMH | 131 | -0.352 | 0.088 | $6.63 \times 10^{-5}$ |
| | WMHicv | 127 | -0.593 | 0.128 | $3.77 \times 10^{-6}$ |
|  | WMicv | 118 | -5929.059 | 3680.657 | 0.107 |
| <u>Social deprivation</u> | gFA | 1 | 0.018 | 0.194 | 0.925 |
|  | gICVF | 1 | -0.135 | 0.190 | 0.478 |
|  | gISOVF | 2 | 0.006 | 0.118 | 0.958 |
|  | GM | 3 | 2392.843 | 1920.525 | 0.213 |
|  | gMD | 3 | 0.202 | 0.097 | 0.038 |
|  | GMicv | 2 | -5209.837 | 6602.841 | 0.430 |
|  | gOD | 2 | -0.133 | 0.126 | 0.288 |

**Table 3.** Showing the IVW causal effect estimate of each indicator of SES on brain measures.
|  |  |  |  |  |
| --- | --- | --- | --- | --- |
| NAWM | 3 | -927.291 | 3898.798 | 0.812 |
| TBV | 2 | 23273.747 | 11127.260 | 0.036 |
| TBVicv | 2 | -8374.616 | 7500.461 | 0.264 |
| WMH | 2 | 0.167 | 0.162 | 0.302 |
| WMHicv | 2 | -0.014 | 0.182 | 0.937 |
| WMicv | 3 | -487.951 | 5232.076 | 0.926 |

For occupational prestige the IVW method indicated a protective causal effect on WMHicv of β = −0.012, SE = 0.006, P = 0.041 (**Table 3, Supplementary Table 7 & Supplementary Figure 8-9**). However, following control for the horizontal pleiotropy identified (Egger intercept = 0.017, SE = 0.008, P = 0.043), this causal estimate increased to β = −0.069, SE = 0.028, P = 0.017 (**Supplementary Table 7**). Horizontal pleiotropy was further examined using MR-PRESSO but no SNPs were detected as outliers (**Supplementary Table 8)**.

The causal effect of education on WMHicv was replicated using an independent sample from the SSGAC. The education replication data set yielded a protective effect against WMHicv of β = −0.186, SE = 0.083, P = 0.026, with no evidence of horizontal pleiotropy, nor was there evidence of heterogeneous effects (Q = 36.138, Q df = 49, Q P = 0.914, **Supplementary Table 9** & **Supplementary Table 10**).

In addition to causal effects on WMHicv, educational attainment was found to have causal effects on WMH (**Table 3** & **Supplementary Table 7)**, where IVW regression showed an effect of β = −0.352, SE = 0.088, P = 6.63×10^−5^. This effect was also replicated (β = −0.140, SE = 0.056, P = 0.012) in an independent sample (**Supplementary Table 9**) with no evidence that it was driven by horizontal pleiotropy (**Supplementary Table 10**).

Despite the lower power of the social deprivation data set both a greater TBV (β = 23273, SE = 11127, P = 0.036) and gMD (β = 0.202, SE = 0.097, P = 0.038) were identified as being a consequence of a greater level of social deprivation (**Table 3, Supplementary Table 7 & Supplementary Figure 10-11**). No other causal effects of SES on brain structure were identified.

### Estimating causal effects of brain structure on indices of SES

In contrast with the causal effects of greater SES on brain imaging traits which showed evidence of causal effects on lower white matter and white matter hyperintensity traits, the causal effects of brain structure on measures of SES provided evidence of total brain volume causally contributing to each indicator of SES: occupational prestige (β = 2.15×10^−5^, P = 4.57×10^−13^), household income (β = 1.67×10^−6^, P = 1.02×10^−22^), and educational attainment (β = 6.50×10^−7^, P = 1.62×10^−13^), and social deprivation (β = −1.84×10^−7^, P = 3.41×10^−8^, **Table 4 & Supplementary Table 7**). MR-Egger regression indicated little evidence of horizontal pleiotropy as the MR-Egger intercept was indistinguishable from zero in each comparison (**Supplementary Table 7)** and MR-PRESSO did not detect any outliers influencing the causal estimate through horizontal pleiotropy (**Supplementary Table 8 & Supplementary Figure 12-14**).

**Table 4.** Showing the IVW causal effect estimate of each indicator of brain structure on socio-economic status. Beta weights are unstandardized and reflect the original unit of measure. Abbreviations: TBV, total brain volume; GM, grey matter volume; WMH, white matter hyperintensity volume; TBVicv, TBV as a proportion of intracranial volume; GMicv, GM as a proportion of intracranial volume; WMicv, white matter volume as a proportion of intracranial volume; WMHicv, WMH as a proportion of intracranial volume; gFA, a general factor of brain white matter tract fractional anisotropy; gMD, a general factor of brain white matter tract mean diffusivity; gIVCF, a general factor of brain white matter tract intracellular volume fraction; gISOVF, a general factor of brain white matter tract isotropic volume fraction; gOD, a general factor of brain white matter tract orientation dispersion.

| **Exposure** | **Outcome** | **N SNPs** | **Beta** | **SE** | **P** |
| --- | --- | --- | --- | --- | --- |
| <u>gFA</u> | Occupation | 34 | 0.156 | 0.251 | 0.534 |
|  | Income | 34 | -0.004 | 0.014 | 0.747 |
|  | Education | 34 | 0.007 | 0.007 | 0.314 |
|  | Social deprivation | 34 | -0.039 | 0.034 | 0.243 |
| <u>gICVF</u> | Occupation | 53 | 0.058 | 0.161 | 0.720 |
|  | Income | 53 | 0.008 | 0.012 | 0.517 |
|  | Education | 53 | 0.003 | 0.005 | 0.586 |
|  | Social deprivation | 53 | -0.037 | 0.022 | 0.093 |
| <u>gISOVF</u> | Occupation | 22 | -0.144 | 0.362 | 0.690 |
|  | Income | 22 | 0.002 | 0.019 | 0.911 |
|  | Education | 22 | -0.008 | 0.010 | 0.441 |
|  | Social deprivation | 22 | -0.015 | 0.037 | 0.681 |
| <u>GM</u> | Occupation | 22 | $1.12 \times 10^{-5}$ | $1.81 \times 10^{-5}$ | 0.536 |
| | Income | 22 | $9.11 \times 10^{-7}$ | $1.28 \times 10^{-6}$ | 0.477 |
| | Education | 22 | $3.57 \times 10^{-7}$ | $5.39 \times 10^{-7}$ | 0.509 |
| | Social deprivation | 22 | $5.03 \times 10^{-7}$ | $2.29 \times 10^{-6}$ | 0.826 |
| <u>gMD</u> | Occupation | 29 | -0.156 | 0.271 | 0.563 |
|  | Income | 29 | -0.016 | 0.020 | 0.420 |
|  | Education | 29 | -0.006 | 0.009 | 0.465 |
|  | Social deprivation | 29 | 0.044 | 0.033 | 0.184 |
| <u>GMicv</u> | Occupation | 20 | $-1.44 \times 10^{-5}$ | $6.99 \times 10^{-6}$ | 0.040 |
| | Income | 20 | $9.01 \times 10^{-8}$ | $4.22 \times 10^{-7}$ | 0.831 |
| | Education | 20 | $-3.70 \times 10^{-7}$ | $2.14 \times 10^{-7}$ | 0.085 |
| | Social deprivation | 20 | $-3.96 \times 10^{-7}$ | $9.94 \times 10^{-7}$ | 0.690 |
| <u>gOD</u> | Occupation | 24 | 0.130 | 0.450 | 0.772 |
|  | Income | 24 | 0.014 | 0.034 | 0.668 |
|  | Education | 24 | 0.003 | 0.015 | 0.854 |
|  | Social deprivation | 24 | -0.032 | 0.033 | 0.335 |
| <u>TBV</u> | Occupation | 34 | $2.15 \times 10^{-5}$ | $2.97 \times 10^{-6}$ | $4.57 \times 10^{-13}$ |
| | Income | 34 | $1.61 \times 10^{-6}$ | $1.64 \times 10^{-7}$ | $1.02 \times 10^{-22}$ |
| | Education | 34 | $6.50 \times 10^{-7}$ | $8.82 \times 10^{-8}$ | $1.62 \times 10^{-13}$ |
| | Social deprivation | 34 | $-1.84 \times 10^{-6}$ | $3.34 \times 10^{-7}$ | $3.41 \times 10^{-8}$ |
| TBVicv | Occupation | 29 | $-4.10 \times 10^{-6}$ | $5.38 \times 10^{-6}$ | 0.445 |
| | Income | 29 | $9.76 \times 10^{-8}$ | $3.44 \times 10^{-7}$ | 0.777 |
| | Education | 29 | $-5.67 \times 10^{-8}$ | $1.63 \times 10^{-7}$ | 0.727 |
| | Social deprivation | 29 | $4.58 \times 10^{-7}$ | $5.11 \times 10^{-7}$ | 0.370 |
| WMH | Occupation | 16 | 0.218 | 0.251 | 0.385 |
|  | Income | 16 | 0.014 | 0.018 | 0.422 |
|  | Education | 16 | 0.003 | 0.008 | 0.674 |
|  | Social deprivation | 16 | 0.022 | 0.039 | 0.583 |
| WMHicv | Occupation | 18 | -0.334 | 0.290 | 0.250 |
|  | Income | 18 | -0.033 | 0.022 | 0.146 |
|  | Education | 18 | -0.011 | 0.009 | 0.246 |
|  | Social deprivation | 18 | 0.067 | 0.035 | 0.055 |
| WMicv | Occupation | 28 | $7.92 \times 10^{-6}$ | $1.07 \times 10^{-5}$ | 0.461 |
| | Income | 28 | $1.28 \times 10^{-6}$ | $5.94 \times 10^{-7}$ | 0.031 |
| | Education | 28 | $2.76 \times 10^{-7}$ | $3.14 \times 10^{-7}$ | 0.379 |
| | Social deprivation | 28 | $-2.30 \times 10^{-7}$ | $8.12 \times 10^{-7}$ | 0.776 |

The causal effects of TBV on both educational attainment (β = 1.01×10^−6^, P = 6.14×10^−6^) and household income (β = 1.23×10^−6^, P = 2.16×10^−6^) were replicated in to independent samples (**Supplementary Table 9**). Neither of these replications showed evidence of horizontal pleiotropy (**Supplementary Table 9**) however MR-PRESSO indicated that that three SNPs were distorting the causal estimate (**Supplementary Table 10**). Following their removal, the significant causal estimate was smaller but retained statistical significance (β = 7.62×10^−7^, P = 2.00×10^−4^).

In addition to total brain volume both WMicv and GMicv showed evidence of causal effects on income, and occupational prestige respectively. However, MR-PRESSO identified four SNPs distorting the causal estimate of WMicv on household income (**Supplementary Table 8**) and following their removal, little evidence of a causal effect was found. Furthermore, no effect of WMicv on household income was identified using the IVW method performed in our replication sample (**Supplementary Table 8**) indicating that this effect was potentially driven by horizontal pleiotropy or a false positive. Similarly the use of MR-PRESSO removed the causal estimate of GMicv on occupational prestige but no individual SNP contributing to horizontal pleiotropy was identified (**Supplementary Table 8**).

### The role of cognitive ability in the link between SES and brain structure

Genetic associations with SES variables are unlikely to be due to the identified variant exerting a biological effect that contributes directly to the observed differences in SES^11^. Rather, the observed SNP-trait association is more likely due to the effects of vertical pleiotropy where genetic variation contributes towards heritable traits that are themselves responsible for the differences in SES^11,12^. Due to the strong genetic^19^ and phenotypic correlations between measures of SES and cognitive ability, as well as the finding that cognitive ability is one of the likely heritable, causal, traits on the phenotypic pathway between genetic inheritance and income differences^11,30,31^, we examine if the heritable traits captured by GWAS conducted on the four indices of SES show causal effects on brain structure that can be explained by cognitive ability.

### Measures of SES capture a set of heritable traits common to each indicator

First, we use the heterogeneity (Q) statistics derived using our common factor model of socioeconomic status and the GWAS results of each individual indicator of SES to examine if SNP effects on the indicators are better explained as SNP effects that act on a latent factor common to each indicator of SES (**Table 5**). Such evidence would be consistent with the idea that a GWAS conducted on each indicator of SES will capture similar heritable traits. FUMA^32^ was used to derive independent genomic loci in the general factor of SES and in each of the four indictors. A total of 469 independent genomic loci were identified for the general factor of SES, and of these 98 loci showed no overlap with any indicator of SES indicating these loci act on the genetic architecture that is shared between each indicator. Occupational prestige, household income, educational attainment, and social deprivation were found to have 68, 73, 491, and 10 independent loci, respectively. However, only eight loci for occupational prestige, 13 for household income and 143 associated with educational attainment, and four for social deprivation were independent from the general factor of SES indicating that the bulk of the genetic effects for each of these SES traits act on the same underlying heritable trait/s.

**Table 5.**
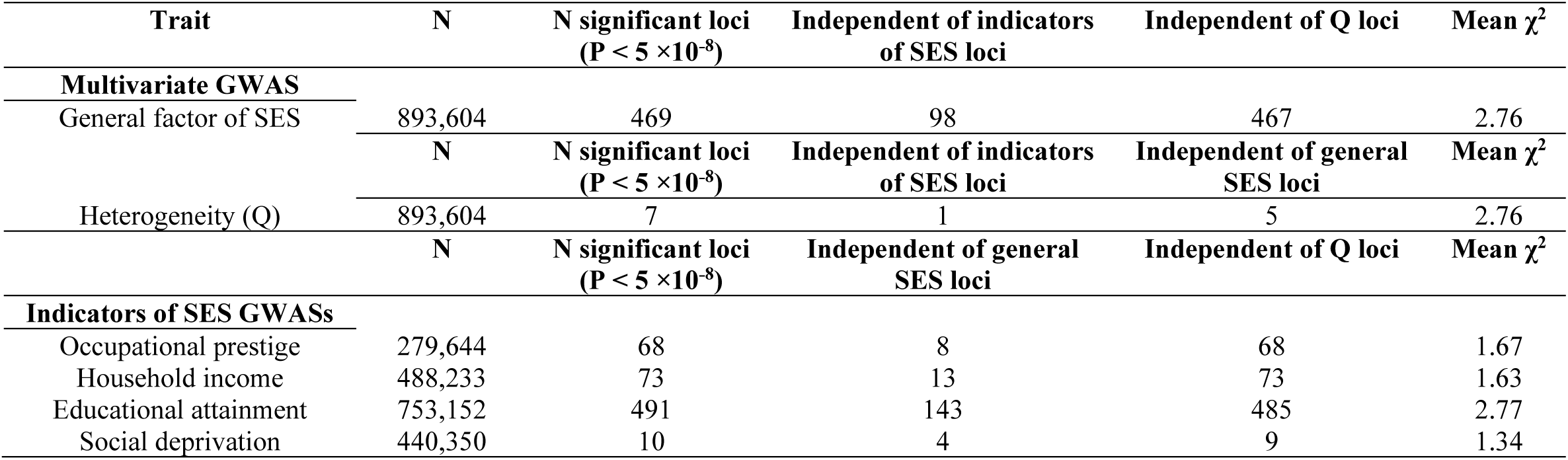
Showing a summary of the general factor of SES multivariate GWAS and the univariate GWAS on each indicator of SES.

| Trait | N | N significant loci<br>($P < 5 \times 10^{-8}$) | Independent of indicators<br>of SES loci | Independent of Q loci | Mean $\chi^2$ |
| --- | --- | --- | --- | --- | --- |
| **Multivariate GWAS** |  |  |  |  |  |
| General factor of SES | 893,604 | 469 | 98 | 467 | 2.76 |
| | N | N significant loci<br>($P < 5 \times 10^{-8}$) | Independent of indicators<br>of SES loci | Independent of general<br>SES loci | Mean $\chi^2$ |
| Heterogeneity (Q) | 893,604 | 7 | 1 | 5 | 2.76 |
| | N | N significant loci<br>($P < 5 \times 10^{-8}$) | Independent of general<br>SES loci | Independent of Q loci | Mean $\chi^2$ |
| **Indicators of SES GWASs** |  |  |  |  |  |
| Occupational prestige | 279,644 | 68 | 8 | 68 | 1.67 |
| Household income | 488,233 | 73 | 13 | 73 | 1.63 |
| Educational attainment | 753,152 | 491 | 143 | 485 | 2.77 |
| Social deprivation | 440,350 | 10 | 4 | 9 | 1.34 |

### Overlap of causal loci for cognitive ability and SES

Using MiXeR^33^, we examined the degree of polygenic overlap between cognitive ability with the general factor of SES, as well as with each indicator of SES. We found that most of the loci associated with each index of SES overlap heavily with loci associated with cognitive ability (**Figure 2**). The genetic relationship between cognitive ability and SES changed, modestly, depending on the measure of SES used, and was not predicted by the genetic correlations. For example, both education and occupational prestige showed a genetic correlation with cognitive ability of ∼ *r_g_* = 0.75, however where education had a large number of education specific causal variants (∼1,500, SE = 700), occupational prestige did not (**Figure 2A**). Furthermore, occupational prestige, which had a genetic correlation with cognitive ability of a similar magnitude as educational attainment (*r_g_* = 0.60), similarly showed no evidence of causal loci that were not also associated with differences on cognitive ability test scores. However, the comparison between occupational prestige and cognitive ability did tentatively indicate that there were loci causal in cognitive ability test score differences that were unrelated to differences in occupational prestige (**Figure 2A & Supplementary Figure 15**). There was little evidence of cognitive ability loci that were not also associated with SES.

**Figure 2.**
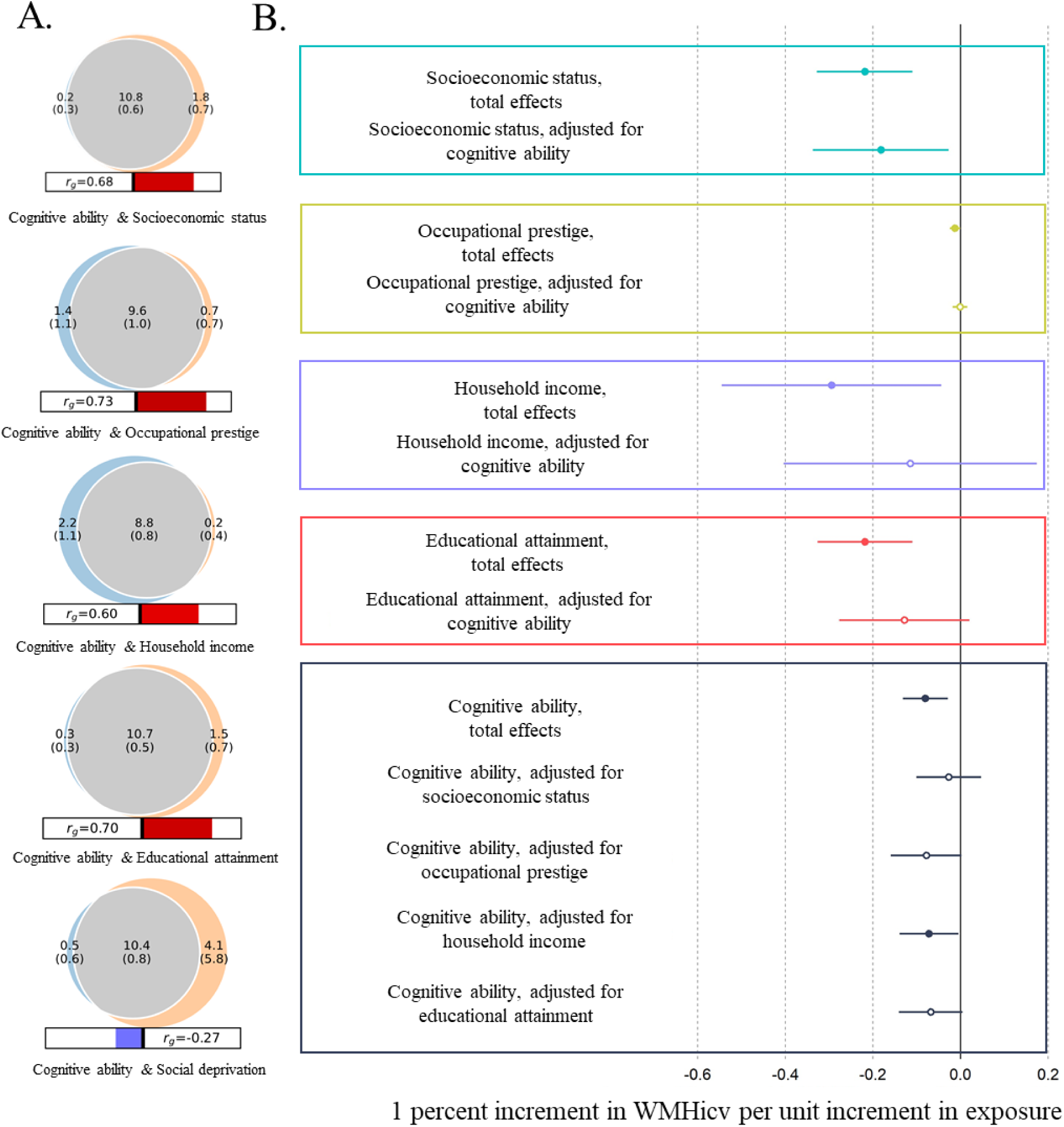
**A.** Venn diagram of cognitive ability and the four indices of SES showing the unique and shared genetic components at the causal level. Grey illustrates the polygenic overlap between trait pairs, orange shows the SES specific components, and blue the unique contributors to cognitive ability. Numbers indicate the estimated quantity of causal variants in thousands with the standard error in brackets. The size of the circle indicates the degree of polygenicity for each trait pair. **B.** Illustrating the total and direct effects of socioeconomic status, occupational prestige, household income, and educational attainment. Colour represents trait and solid shapes indicate a statistically significant causal estimate. Error bars show ± one standard error

### Estimating the bidirectional effects between cognitive ability and socioeconomic status

Previous work has indicated a bi-directional causal relationship between educational attainment with cognitive ability^30,31^ and that cognitive ability is one of the causal factors in income differences^11^. Here we show a bi-directional causal relationship between the general factor of SES with cognitive ability, and a bi-directional causal relationship between each indicator of SES with cognitive ability (**Table 6** & **Supplementary Table 11**). Following Steiger filtering, we find evidence that higher cognitive ability was causally linked to having a higher level of SES using the general factor of SES (β =0.192, SE = 0.012, P = 1.24×10^−53^). Cognitive ability was also a causal factor in each specific indicator of SES and was linked with a higher level of occupational prestige (β = 2.67, SE = 0.15, P = 2.21×10^−69^), a greater level of household income (β = 0.131, SE = 0.011, P = 2.66×10^−31^), a greater chance of attaining a university level education (β = 0.077, SE = 0.004, P = 5.75×10^−67^), and decrease in level of deprivation in which one lives (β = −0.84, SE = 0.025, P = 0.001, **Table 6** & **Supplementary Table 11**). There was evidence of heterogeneity in each estimate of the causal effects of cognitive ability on SES as indicated by significant Cochran’s Q statistics^28^ (**Supplementary Table 11**). This heterogeneity statistic provides an indication of the variability of the estimated effect between SNPs and can arise if the SNPs have horizontal pleiotropic effects. However, there was little evidence that horizontal pleiotropy biased the causal estimates of cognitive ability on SES; the MR Egger regression intercepts were close to zero and MR PRESSO indicated no significant distortion in the causal estimate due to SNPs with a horizontal pleiotropic effect (**Supplementary Table 12, & Supplementary Figures 16-18**).

**Table 6.**
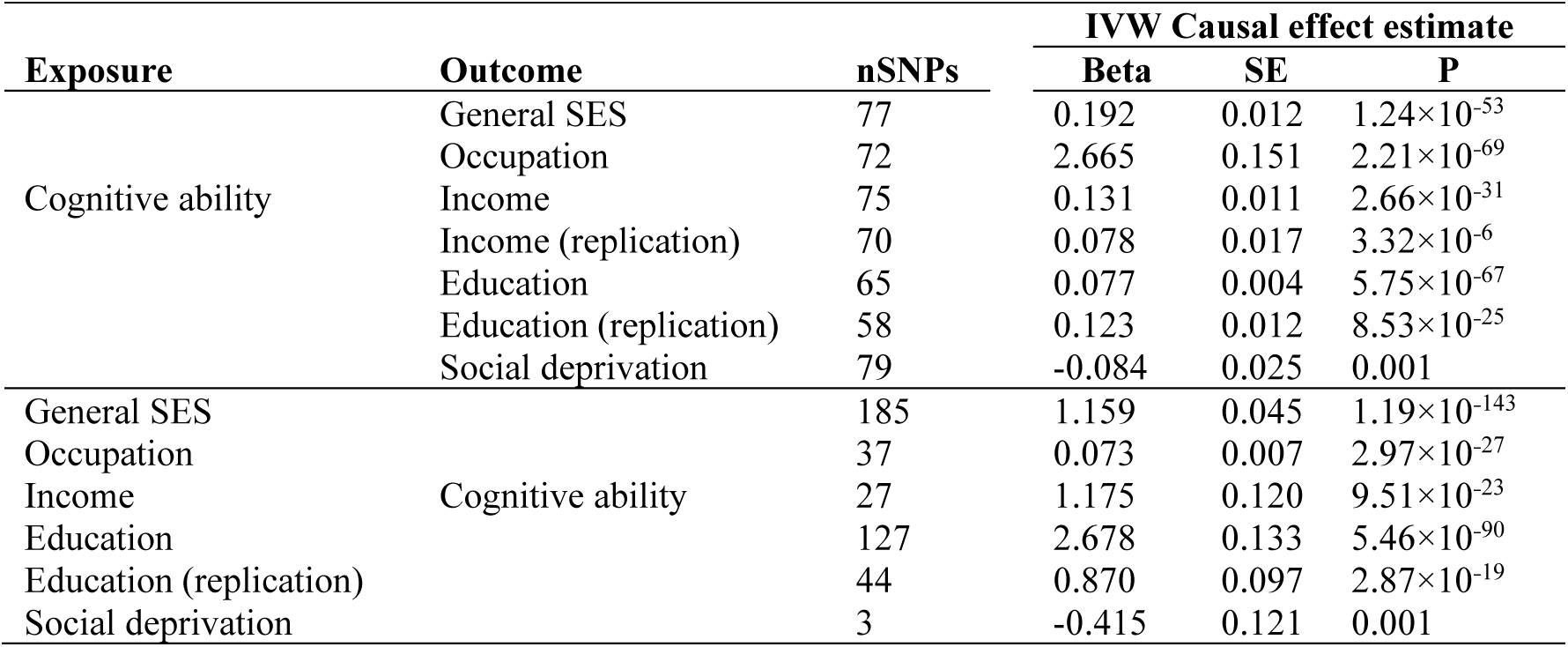
Showing the bi-directional total causal effects of cognitive ability on the general factor of SES and each of the four indicators of SES. Beta weights are unstandardized and reflect the original unit of measure.

Using the GWASs conducted on each SES variable, instrumental variables were identified to examine if they captured heritable traits that were causal factors in differences in cognitive ability. We find evidence that in addition to the causal effect on the general factor of SES (β = 1.159, SE = 0.045, P = 1.19×10^−143^), increases in cognitive ability were a consequence of increases in education (β = 2.433, SE = 0.129, P = 7.54×10^−90^), income (β = 1.103, SE = 0. 116, P = 2.17×10^−21^), and occupational prestige (β = 0.071, SE = 0.007, P = 7.93×10^−27^), whereas there was weak evidence that the heritable traits linked to increases in social deprivation acted to reduce cognitive ability (β = −0.415, SE = 0.121, P = 0.001, **Supplementary Table 11**). As with the causal effects of cognitive ability on SES there was significant heterogeneity in the estimates (**Supplementary Table 11**) but little evidence of bias arising due to horizontal pleiotropy indicated by the MR Egger intercepts not being significantly different from zero and no distortion detected using MR-PRESSO (**Supplementary Table 12** & **Supplementary Figures 19-21**).

### The bidirectional causal effect of cognitive ability on brain structure

Consistent with the idea that GWASs of SES capture variance in cognitive ability, we find evidence that cognitive ability has a protective causal effect on WHMicv (β = −0.080, SE = 0.026, P = 0.002). Furthermore, and again consistent with what was identified for measures of SES, there was evidence to suggest a greater total brain volume resulted in a higher level of cognitive ability (β = 3.97×10^-6^, 4.96×10^-7^, 1.28×10^-15^). No evidence of horizontal pleiotropy was identified using MR Egger (Egger_intercept_ P = 0.492) and MR-PRESSO found no evidence of distortion in the causal estimate following the removal of five SNPs with evidence of horizontal pleiotropy (MR-PRESSO distortion P-value = 0.516). There was however, significant heterogeneity in the causal estimate of TBV on cognitive ability (Q P-value = 1.85×10^-10^, **Supplementary Tables 13-14** & **Supplementary Figures 22-24**).

### Direct effects of SES on brain structure

Using Multivariable MR (MVMR)^34^ to control for the effects of cognitive ability on each of the SES variables we examine if the causal effects of each SES variable on brain structures were independent of cognitive ability.

When both cognitive ability and the general factor of SES were included in a single multivariate model there was evidence that SES effects on WMHicv that were independent of cognitive ability (total effect β = −0.218, SE = 0.056, P = 8.63×10^−5^, direct effect β = −0.182, SE = 0.079, P = 0.022).

However, for the indicators of SES, following adjustment for cognitive ability, a steep decline of in the point estimate of the effect sizes was evident when comparing between the total and direct effects (**Figure 2B**). Furthermore, beyond the effects of cognitive ability on WMHicv there were no direct effects of occupational prestige (total effect β = −0.069, SE = 0.028, P = 0.017, direct effect β = −0.001, SE = 0.008, P = 0.886, 98% reduction of direct effect), household income (total effect β = −0.294, SE = 0.128, P = 0.021, direct effect β = −0.115, SE = 0.147, P = 0.436, 61% reduction of total effect) or educational attainment (total effect β = −0.218, SE = 0.055, P = 7.64×10^−5^, direct effect β = −0.128, SE = 0.076, P = 0.091, 41% reduction of total effect, **Figure 2B**).

The reduction in the causal estimate of cognitive ability on WMHicv following the inclusion of each indicator of SES was more modest (range of reduction 15%-2%) with cognitive ability still retaining a significant causal effect on WMHicv following the inclusion of household income (total effect β = −0.08, SE = 0.026, P = 0.002, direct effect β = −0.078, SE = 0.034, P = 0.036). Following the inclusion of occupational prestige (direct effect β = −0.078, SE = 0.041, P = 0.057), educational attainment (direct effect β = −0.068, SE = 0.037, P = 0.068), of the general factor of SES (direct effect β = −0.027, SE = 0.038, P = 0.48) there was no significant effect of cognitive ability on WMHicv (**Supplementary Table 15**).

## Discussion

Those individuals from more advantaged socioeconomic backgrounds will typically have fewer instances of poor physical and mental health compared to those from more deprived backgrounds^1,5–7^. Understanding the causes of such differences has the potential to decrease health disparities and improve our understanding of the intricate working of societal risk factors of illnesses. In the current study we examine the role that SES plays on brain structure by performing a multivariate GWAS to capture sources of SES differences that effect the individual, the household, and the area in which one lives. Our GWAS on SES was then used to derive instrumental variables to examine the causal effect differences in SES has on brain morphology and health. The current study contributes to our understanding of the genetic contributions to SES in at least five ways.

First, we show that whilst a common phenotypic factor explains only 31.2% of phenotypic variation across each indicator of SES, our multivariate general genetic factor of SES accounted for on average 76% (range =49%-93%) of the genetic variation found across occupational prestige, household income, educational attainment, and social deprivation. Furthermore, our common factor model showed that the same heritable traits underlie the bulk of the heritable variation in SES across each of the indicators where, of the 469 independent genomic loci identified, only two showed evidence of a heterogenous effect indicated by a significant Q value. This asymmetry in the variance captured by a common phenotypic factor of SES compared with the variance captured by a common genetic factor of SES, and the finding that the majority of loci associated with the general factor acted on each of the four indicators of SES, implies that although each indicator captures a different environmental component of SES, the heritable traits that give rise to these phenotypic differences are largely the same.

The identification of this common genetic factor of SES allows for the recontextualisation of the results of previous GWAS that have been conducted on individual indicators of SES. Specifically, many of the loci identified in univariate GWAS of a single indicator of SES are generalisable to SES more broadly, as they are associated with all indicators that load on the general genetic factor of SES. For example, previous GWAS examining educational attainment^17^ and income^11^ have reported 3,952 and 149 loci respectively as showing an association with a specific indicator of SES. Here, we find that 78.8% of the genetic variance of educational attainment and 84.2% of the genetic variance of income is through this general factor of SES indicating that only a minority of the loci captured by those GWAS on specific indicators of SES will be trait specific.

Second, we find evidence that cognitive ability is one of the likely causal traits captured by GWAS on SES. By using MiXeR^33^ we show that of the estimated 11,000 causal variants for cognitive ability, 10,800 are shared with the general factor of SES with only 1,800 causal variants for SES not shared with cognitive ability. Whilst MiXeR cannot differentiate between vertical and horizontal pleiotropy^33^, across each indicator of SES there was little evidence of loci associated with cognitive ability that were not also associated with differences in SES, consistent with the hypothesis that differences in cognitive ability are one of multiple heritable traits that influence differences in SES.

By using Two-sample MR we were able to confirm that vertical pleiotropy, and not horizontal pleiotropy, best explained the overlapping genetic architecture between cognitive ability and SES identified using MiXeR. Higher cognitive ability was one of the causal elements of having a greater level of the general factor of SES, a higher occupational prestige and educational attainment, a higher household income, and living in a less deprived environment. This effect was replicated using educational attainment and household income data sets that included participants from outside the UK indicating these effects were not specific to the UK or to the participants of UK Biobank. These effects were bidirectional and differences in SES were also shown to influence cognitive ability.

Third, using Two-sample MR we show that higher levels of this common factor of SES is a consequence of a greater total brain volume and a likely causal factor in lower levels of white matter hyperintensities (WMHicv). White matter hyperintensities are white matter lesions that, on fluid attenuated inversion recovery (FLAIR) MRI scans, show a signal intensity that is brighter than surrounding white matter^35^. WMHs are associated with vascular risk and small vessel disease^36^ and may indicate permeability in the blood brain barrier as well as axonal and myelin degeneration^37^ Furthermore, increases in WMH volume are associated with cognitive decline and higher risk of Alzheimer’s disease, as well as with lower levels of cognitive ability^38^.

In the context of non-clinical community-dwelling adults, WMH volume is also a frank marker of neurodegeneration, being of extremely low prevalence in young adulthood^39^. However, lower levels of cognitive ability at age 11 are associated with greater WMH volume at age 73^40^ indicating that they may influence the trajectory of cognitive decline in adulthood and older age. Our finding that SES was a likely causal factor for WMHicv indicates that lower levels of SES act as a risk factor for the development of WMH across the adult lifespan and may, through the accumulation of damage caused by WMH, increase the rate of cognitive decline and the likelihood of a dementia diagnosis in older age. In contrast, our finding that TBV was a causal factor for SES and cognitive ability may indicate that TBV (which reaches its peak in early adulthood^41^) is a risk factor that influences the rate of cognitive development in childhood.

Fourth, we show using MVMR, that there are causal effects of SES on WMHicv independent of cognitive ability. In the same way a polygenic score captures the aggregate effect of the SNPs used in its construction^42^, so each SNP in a GWAS conducted on SES will capture the aggregate effect of each heritable trait linked to differences in SES^12^. Using MVMR we were able to remove the effect of one of these traits, cognitive ability, in order to gauge the effect of the remaining traits captured by SES on brain morphology. In doing so we show that the direct effects of SES are protective against WMHicv. This is consistent with the idea that the general factor of SES captures a constellation of risk from multiple genetically influenced traits and higher levels of SES are not protective solely due to them capturing differences in cognitive ability^10,11^. These traits could be social health factors^43^ or aspects of personality linked to health, such as conscientiousness, which is phenotypically linked with lower instance of disease^44^ and greater longevity^45^ and shows genetic correlations with mental health traits such as MDD, ADHD, and schizophrenia^46^. Previous work examining educational attainment, an indicator of SES, identified that differences in wellbeing, health, and personality have been shown to make a contribution to the heritability of educational attainment that is independent from cognitive ability^10^.

Fifth, with the qualified exception of educational attainment, which showed ∼15,000 causal loci not also linked to differences in cognitive ability, no evidence was found for loci associated with indicators of SES that were not also loci associated with cognitive ability. However, we find no evidence that these non-cognitive aspects of educational attainment^47^ were causally associated with WMHicv.

Our study has limitations that should be considered when interpreting the results. First, all samples used were from western European societies and cultures of the 21^st^ century. The importance of this caveat is underscored by the observation that the heritable traits that give rise to differences in SES are unlikely to be universal and will be specific to the cultures and samples examined^13,14^. Without studies aiming to examine the heritable traits that give rise to SES and the role these play in brain structure in other cultures, meaningful comparisons between the present study and other cultures are unwarranted.

Second, genetic variants captured by our measures of SES are likely to have pleiotropic effects^48^. To satisfy the assumptions that the genetic association with the outcome is entirely mediated via the exposure, we performed Steiger filtering to remove variants that are more strongly associated with outcome than the exposure (i.e. reverse causation). Although removing invalid instrumental variables and only keep likely vertical pleotropic instrumental variables can improve the validity of causal effects, such data-driven selection of instrumental variables may yield over precise causal effects, especially when the majority of instrumental variables are affected by heterogeneity. Furthermore, in order to break the assumptions of MR it is not sufficient for the genetic variants in the instrumental variable to have pleiotropic effects^49^, rather the genetic variants must have horizontally pleiotropic effects that are mediated through mechanisms other than those captured by SES. For example, should genetic variants have vertically pleiotropic effects, e.g. SNP->neuron-> cognitive ability −>education->income->health->brain structure, then our MR derived causal estimates will not be biased. Furthermore, should the SNPs affect other phenotypes, but these phenotypes do not affect the outcomes, then our MR estimates will not be biased. Whilst it is possible that the genetic variants identified in our GWAS conducted on measures of SES do have horizontally pleiotropic effects, it is unclear what mechanisms would mediate such effects (e.g. personality). In the current study we investigate potentially pleiotropic effects using multivariable Mendelian randomization to examine the role of cognitive ability. Future research should use multivariable Mendelian randomization to investigate this the role of other traits that link SES to brain structure.

Third, there is the potential that indirect genetic effects will contribute to the MR estimates^50^. Indirect genetic effects refer to one individual’s genotype influencing the outcome of another individual’s phenotype, for example, a parent providing material resources for their offspring which may affect SES or cognitive ability. Detecting the magnitude of potential bias resulting from dynastic effects is challenging outside of using family-based data, and at present no such data exist.

Finally, molecular genetic studies examining traits such as cognitive ability and socioeconomic status are prone to misunderstanding and mischaracterisation. These mischaracterisations can include arguments based around genetic determinism where the role of the environment is disregarded in favour of creating myths about immutable, biological differences underlying trait variation, something incompatible with current knowledge of complex traits. In order to communicate our research findings to a general reader in an ethical and socially responsible way, we have provided an FAQ document in **Supplementary Note 1**.

Overall, this study offers new insights into the complex interactions between socioeconomic status (SES), brain development and the risk factors underlying cognitive decline. Employing modern analytical methods on extensive datasets, the findings significantly contribute to our comprehension of factors that influence physical and mental health. Ultimately, these results could highlight potential modifiable risk factors for maintaining cognitive ability in older-age.

## Methods

### Samples

European samples from UK Biobank^51^ were retained if they had genetic information available, sex that was consistent between self-reported and inferred using genotype, no sex chromosome aneuploidies, not having been detected as extreme outliers of heterozygosity and missingness, having not withdrawn consent, and having a genotyping rate greater than 0.9. This resulted in 435,340 participants being available for analysis. European ancestry was identified from the UK Biobank participants that self-reported as white. Principal components (PC) were derived from the genotype data and participants were excluded if they were outside of a mean ± 3 standard deviations from the first six principal components. For our general factor of SES we used all participants who had provided phenotypic data on at least one of our measures of SES.

For our Mendelian randomisation analysis we derived two independent samples using the participants of UK Biobank. The brain imaging subset which consisted of 38,371 participants that had at least one MRI phenotype, and the SES and cognitive ability group that consisted of 383,220 participants who did not have any MRI phenotype and were not genetically related to anyone in the outcome set based on the pairwise kinship reported by UK Biobank. Ethical approval was granted by UK Biobank and this project was conducted under UK Biobank application 10279.

### Exposures and outcomes

Two-sample Mendelian randomisation (MR) was used to examine the causal effects of cognitive ability and socioeconomic status on 13 structural brain imaging measures.

Five cognitive and socioeconomic status variables were considered from UK Biobank, cognitive ability, income, social deprivation, occupational prestige, and educational attainment. Income was measured at the level of the household (HI, *N*=327,402), which was measured in UK Biobank using an ordinal scale of 1 – 5 corresponding to the participants level of household income before tax (1 = < £18,000, 2 = £18,000 −£30,999, 3 = £31,000 − £51,999, 4 = £52,000 - £100,000, 5 = >£100,000).

Social deprivation was measured using the Townsend deprivation index (TS, *N*=382,030). The Townsend deprivation index is an area-based measure of SES derived using the participant’s postcode. Townsend scores were calculated immediately prior to joining UK Biobank and are formed from four measures: the percentage of those aged 16 or over who are unemployed, the percentage of households who do not own a car, do not own their own home, and which are overcrowded. Scores were multiplied by −1 when used for deriving phenotypic and genetic correlations as well as for use in in Genomic SEM to ensure that the direction of effect was the same across each measure of SES (i.e. a greater score indicates a higher level of SES). However, for use in Mendelian randomisation the original direction of effect is retained where a greater score indicates higher level of deprivation (i.e. a lower level of SES).

Occupational prestige was measured using the Cambridge Social Interaction and Stratification Scale (CAMSIS, *N*=242,776) and was derived using job code at visit (data field 20277) in UK Biobank transformed using the method described by Akimova et al. (2023)^52^. In brief, the CAMSIS uses the idea social stratification acts to create differential association where partners and friends are typically selected from within the same social group. Thus, CAMSIS captures the distance between occupations by measuring the frequency of social interactions between them.

Educational attainment (EA, *N*=377,477) was measured by transforming educational qualifications found in UK Biobank to a binary variable where ‘1’ indicated that the participant had attained a university level degree and ‘0’ indicated that they had not.

Due to the high genetic correlations between measures of SES and cognitive ability^17,19^ and the finding that cognitive ability is a likely causal variable in differences in SES in the UK^11,30,31^ cognitive ability was also included as an exposure variable. Cognitive ability was measured using the verbal-numerical reasoning test (VNR, N = 183,321) in UK Biobank. This test consists of 13 (14 for the online version of the test) multiple-choice questions (six verbal and seven numerical) which are to be completed within a two-minute time limit. A participant’s score on each of the questions is then summed to provide an overall measure of the participant’s level of cognitive ability. Participants either completed the VNR test at the assessment centre at one of four time points or completed an online version of the VNR test. If participants took the VNR at multiple time points, only the first instance of the test was used to avoid capturing practise effects in the assessment of the participant’s level of cognitive ability.

Brain structural and diffusion neuroimaging data were acquired, processed and QCd by the UK Biobank team as Imaging Derived Phenotypes (IDPs) according to open access publications^53,54^ and online documentation (https://biobank.ctsu.ox.ac.uk/crystal/crystal/docs/brain_mri.pdf). Global macrostructural outcomes of interest were: total brain volume (TBV), total brain volume as a proportion of intracranial volume (TBVicv), total grey matter volume (GM), total grey matter volume as a proportion of intracranial volume, (GMicv), white matter hyperintensity (WMH) volume, white matter hyperintensity volume as a proportion of intracranial volume (WMHicv), normal-appearing white matter volume (NAWM, total white matter volume – WMH), white matter volume as a proportion of intracranial volume (WMicv), and five global white matter microstructural measures.

The latter were derived from twenty-seven major white matter tracts, for which five tract-averaged white matter diffusion properties were available as IDPs (UK Biobank Category ID 135): fractional anisotropy (FA), mean diffusivity (MD), intra-cellular volume fraction (ICVF), isotropic volume fraction (ISOVF) and orientation dispersion (OD). We ran five PCAs of all 27 tracts; a separate model for each of the five properties. The first unrotated component of each PCA was extracted for further analysis, yielding five global white matter measures (gFA, gMD, gICVF, gISOVF and gOD) which explained 44%, 50%, 68%, 37% and 26% of the variance, respectively. Prior to analysis, participants with the following conditions (UK Biobank field ID 20002.2) were excluded at the outset: dementia, Parkinson’s disease, Guillain-Barré, multiple sclerosis, stroke, brain haemorrhage, brain/intracranial abscess, cerebral aneurysm, cerebral palsy, encephalitis, epilepsy, head injury, infection of the nervous system, ischaemic stroke, meningioma, meningitis, motor neurone disease, spina bifida, subdural haematoma, subarachnoid haemorrhage, transient ischaemic attack, brain cancer, meningeal cancer, other demyelinating or other chronic / neurodegenerative illness, or other neurological injury/trauma. Outliers (>4SDs from the mean, which represented <0.1% of the data in all cases) were then removed from all IDPs prior to analyses. As detailed above, there was no sample overlap between the participants who provided brain imaging data and the participants who provided data pertaining to their SES or cognitive ability.

### Study design and data sets

A valid inference from MR is dependent on satisfying three assumptions: relevance, meaning that the genetic variants must be associated with the risk factor of interest; independence, that the there are no unmeasured confounds of the associations between genetic variants and the outcome; exclusion restriction, that the genetic variants affect the outcome only through the effect they have on the exposure^55^.

Instruments for each exposure were identified using SNPs that attained genome-wide significance (P < 5×10^-8^). These SNPs were then clumped using the 1000G European reference panel and an r^2^ = 0.001, with a 10 Mb boundary. The most significant SNP in each clump was used as an instrumental variable. As all GWAS conducted for this study were performed on the same strand, no palindromic SNPs were excluded from these analyses. The effect of each SNP on the exposure and on the outcome was harmonised to ensure that the effect allele is the same across the exposure and the outcome traits. Steiger filtering was used to ensure that the detected direction of effect (i.e. from exposure to outcome) was correct. Non-Steiger filtered results are also available in **Supplementary Tables 16-20**.

Inverse variance weighted (IVW) regression was used to examine identify putatively causal effects. If there is only one SNP to be used as an instrumental variable, Wald ratio was used. Sensitivity analyses were conducted using MR Egger regression and MR Pleiotropy Residual Sum and Outlier (MR-PRESSO).

Genetic contributions to SES traits are unlikely to be due to a direct genetic effect and is probably the result of multiple heritable traits^11,12^. Furthermore, cognitive ability shows high genetic correlations with measures of SES^19^ and is likely to be one of the causal heritable traits between genetic inheritance and differences in SES^11^. We applied Multivariable Mendelian Randomisation (MVMR)^34^ to examine the causal effects of SES independent of cognitive ability. For MVMR SNPs that were genome-wide in both exposures were retained. Steiger filtering was applied for both exposures on the outcome.

### Replication data sets

Replication of significant causal effects was examined using independent GWAS data set of educational attainment (measured as the number of years of schooling an individual has completed)^15^ and household income (measured as the total annual household income prior to tax)^56^. Following the removal of the participants of UK Biobank, the sample sizes were educational attainment N= 324,162, and household income N = 108,635 (**Supplementary Table 2**). The replication data set for education showed a large significant genetic correlation of *r_g_* = 0.960, SE = 0.015, P = 0 with education in UK Biobank, as did the two household income data sets *r_g_* = 0.957, SE = 0.065, P = 0.

### Meta analysis of income and education

Data provided by the SSGAC was used to add power to the general factor of SES as well acting as a replication sample for educational attainment and household income and for use in MVMR. For both meta-analyses, METAL^57^ was used to conduct a sample size weighted meta-analysis from which Beta values and standard error obtained using the following equation as provided by Zhu et al. (2016)^58^.

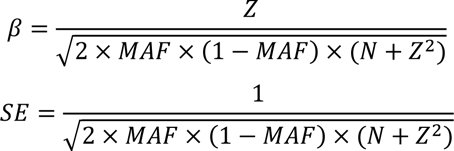

where 𝑀𝐴𝐹 is the minor allele frequency, 𝑁 is the sample size, and 𝑍 is the test-statistics.

### Genome-wide association studies

Genome-wide association studies (GWASs) were conducted in Regeine v3.1.3^59^. Regeine uses a two-step approach to account for sample relatedness and population structure. In the first step, a whole genome regression model was fit to each trait (Exposures and outcomes) using 564,253 genotyped variants. These variants have minor allele frequency (MAF) > 0.01, call rate > 0.9, and Hardy-Weinberg Equilibrium of HWE-p value > 10^-15^.

In the second step, an association test was performed for each of the 13,192,861 imputed variants using a LOCO (leave-one-chromosome out) scheme. These variants have MAF > 0.001 and INFO > 0.8. For binary phenotypes (i.e. Educational attainment), firth logistic regression test was performed in the second step.

The per-chromosome LOCO genomic predictions produced in the first step were fitted in the second step to account for sample relatedness and population structure. In addition, sex, age at assessment, assessment centres, genotyping array, genotyping batch, and the first 40 PCs derived from genotype data were fitted as covariates in both steps. For cognitive ability, participants’ who took the VNR at an assessment centre were analysed together including time point (1-4) as an additional covariate before being meta-analysed with the participants whose first instance of taking the VNR was online. Regarding brain imagining phenotypes, three-dimensional head position along the X, Y, and Z axis were fitted as extra covariates. For TBV height was fitted as an additional covariate and for GM and NAWM both height and TBV were fitted. For VNR, the GWASs were performed in participants who took test in the assessment centre, and those took online test separately, before combining the results with an inverse variance weighted model^60^.

### Linkage Disequilibrium Score Regression (LDSC)

Using the 1000G European reference panel LDSC^24^ was performed to estimate the heritability of the exposure and outcome traits. In addition the intercept of each LDSC regression was used to examine the GWAS association test statistics for inflation due to factors other than polygenicity.

### MiXeR

MiXeR v1.3 (https://github.com/precimed/mixer) was used to examine the genetic overlap between cognitive ability and SES traits. First, a univariate model^61^ was run to study the polygenicity (i.e. number of variants) of each trait using the Z-score from GWAS summary statistics and 1000G European LD panel. Second, a bivariate model^33^ was used to estimate the genetic overlap (i.e. number of variants shared between cognitive ability and SES traits) using the parameters learned from the univariate model. The analysis was repeated twenty times using 2 million randomly selected SNPs at each time. The results across twenty runs were then averaged and the genetic overlap of the best model with the lowest –log likelihood ratio was plotted (**Supplementary Figure 15)**.

### Phenotypic and genomic structural equation modelling

Phenotypic common factor of SES was derived in R^62^ using factor analysis in *psych*^63^ package on standardised occupational prestige (n = 279,644), household income (n = 488,233), educational attainment (n = 753,152), and social deprivation (n = 440,350) phenotypes. Note that as sample overlap is controlled for in Genomic SEM these samples sizes are larger than those used in our Two-sample Mendelian randomisation analysis described above. Regarding genetic common factor of SES, we used genomic structural equation modelling^25^ to derive LDSC—based^48^ genetic correlations and covariances between occupational prestige, household income, educational attainment, and social deprivation. Next, the covariance structure between each of the four measures of SES was used to derive a genomic structural equation model to examine their loading on a single factor of SES. This common factor model was ran using SNPs from each indicator of SES where MAF >0.01 and INFO > 0.9. We then performed a multivariate GWAS using genomic SEM where 7,462,726 SNPs with MAF >0.01 and INFO >0.6 were included to derive genome-wide summary statistics describing each SNPs association with the common factor of SES. In addition, we derived genome-wide heterogeneity (Q) statistics describing the degree to which a given SNP is likely not acting on single latent factor of SES.

### Loci identification and overlap

For each trait, genomic risk loci were identified by FUMA^32^ (version v1.3.6a) using 1000G EUR reference panels. Briefly, FUMA performed two LD clumpings. The first clumping was designed to define independent signals (genome significant SNPs at P < 5×10^-8^) with *r*^2^ > 0.6. In the second clumping, independent signals were clumped into one genomic locus if the *r*^2^ between two signals is > 0.1 or two signals are within 250kb. The SNPs clumped into each genomic locus naturally formed its physical boundary.

We compared the positions of genomic loci between two traits locus-by-locus. We define that a locus of trait A overlaps with trait B, if the positions of any trait B loci overlap with the position of that trait A locus. For the general factor of SES, we define a locus is unique to general SES if that locus does not overlap with any of the four contributing traits. For the four contributing traits of general SES, we define a locus is unique to that trait if that locus does not overlap with general SES.

## Data availability

Summary statistics GWASs for the general factor of socioeconomic status, social deprivation, occupational prestige, and the discovery GWAS data set for household income, and educational attainment will be available on GWAS catalog upon publication (https://www.ebi.ac.uk/gwas/). The replication samples are available on request from the Social Science Genetic Association Consortium (https://www.thessgac.org/).

## Acknowledgements

W.D.H., and C.X. are supported by a Career Development Award from the Medical Research Council (MRC) [MR/T030852/1] for the project titled “From genetic sequence to phenotypic consequence: genetic and environmental links between cognitive ability, socioeconomic position, and health”. For the purpose of open access, the author has applied a ‘Creative Commons Attribution (CC BY) licence to any Author Accepted Manuscript version arising from this submission.

## Author contributions

All authors approved the final manuscript

