## Supplementary Figures for "Deciphering the Influence of Socioeconomic Status on Brain Structure: Insights from Mendelian Randomization"

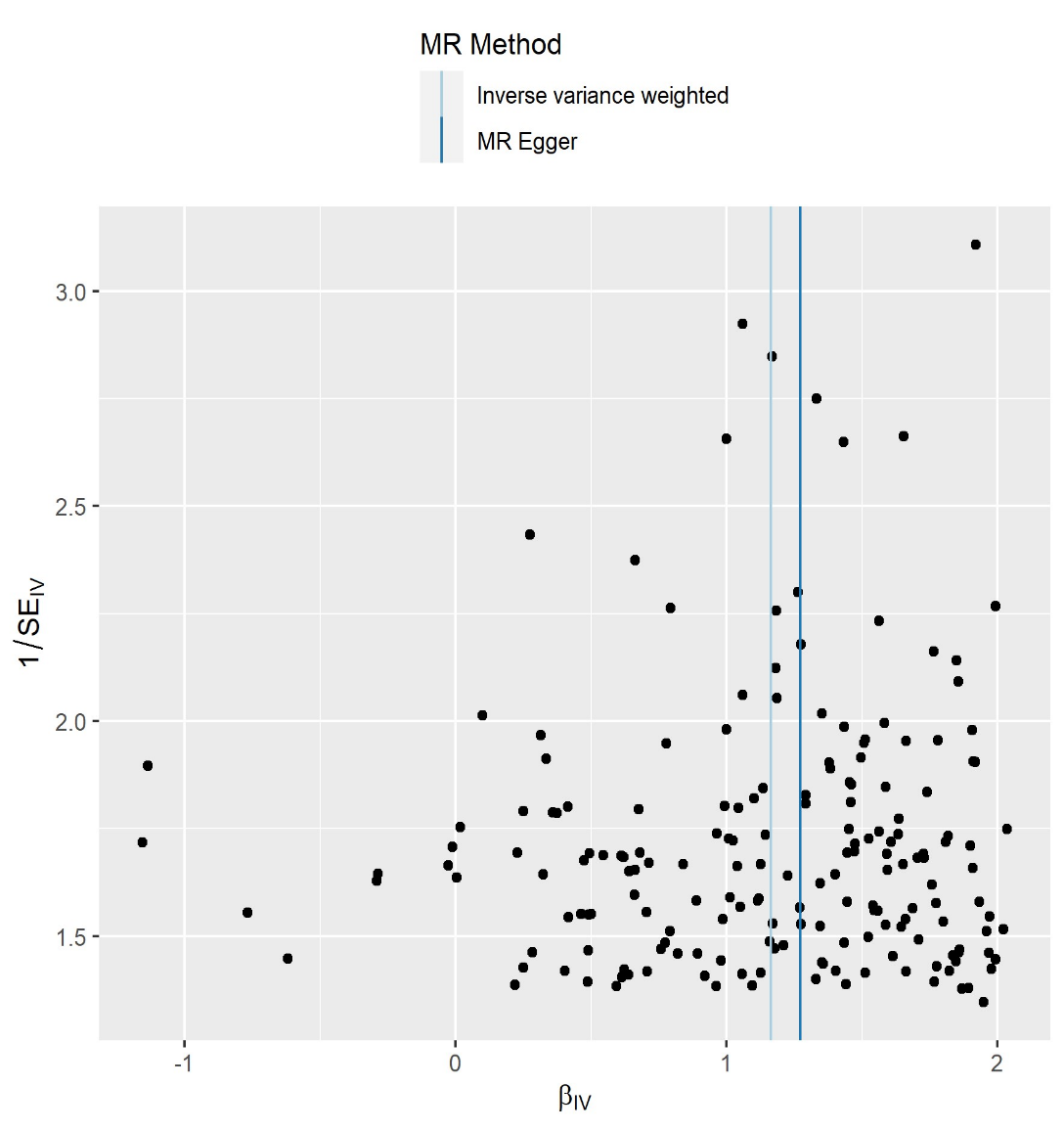

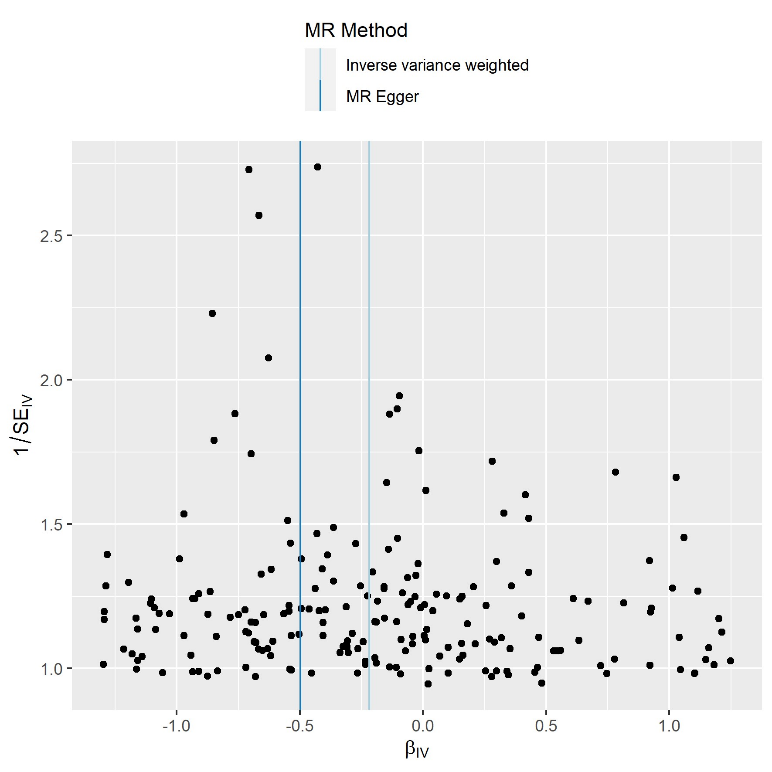

WMHicv

**Supplementary Figure 1.** Funnel plots used to examine the extent to which pleiotropy is balanced across the instruments used in the univariate Mendelian randomisation of general factor of socioeconomic status on white matter as a proportion of intracranial volume and white matter hyperintensities.

### Leave one out plot Socioeconomic status on WMHicv

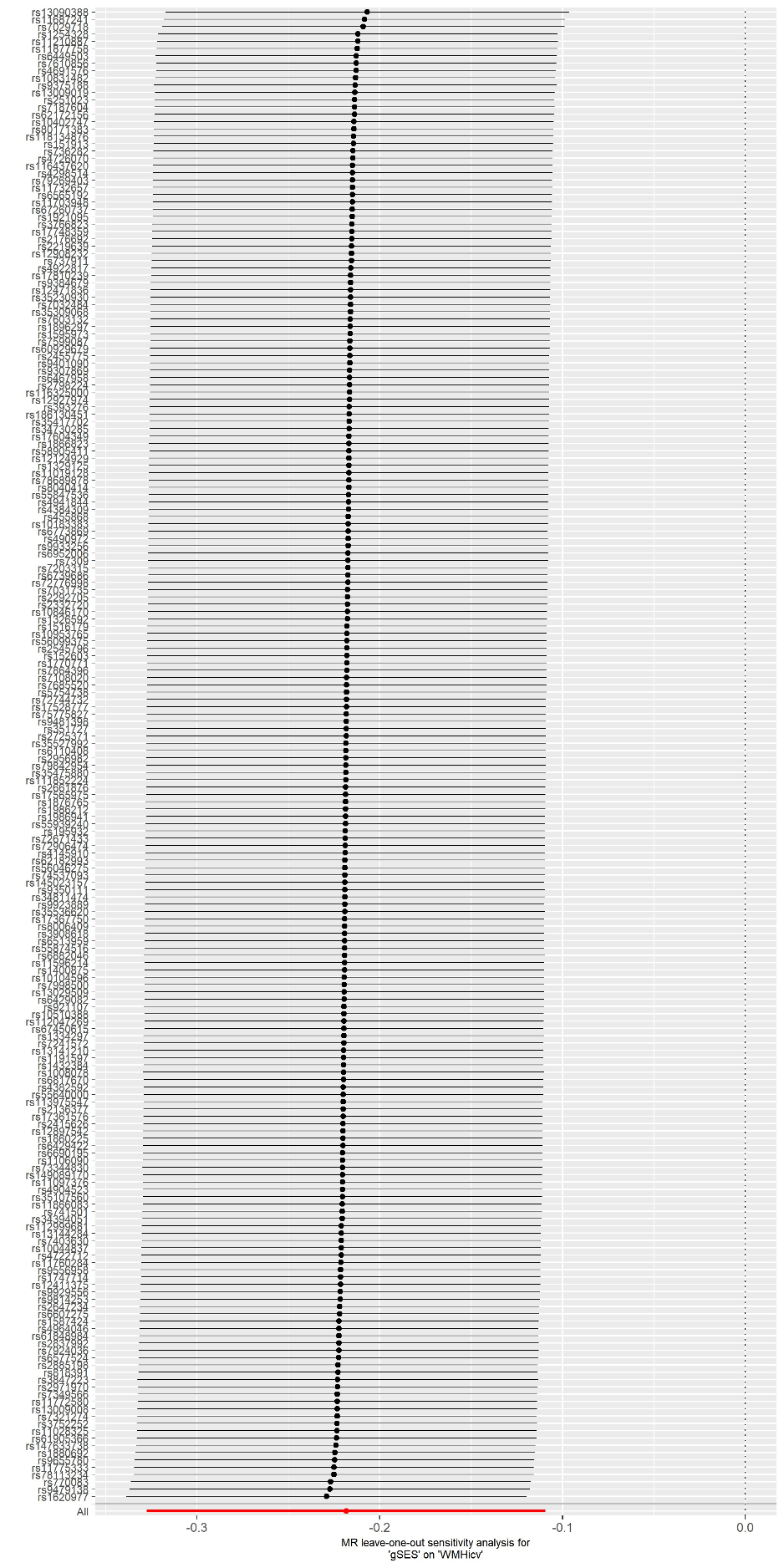

**Supplementary Figure 2.** Leave one out plot of the univariate Mendelian randomisation of general factor of socioeconomic status on white matter hyperintensities as a proportion of intracranial volume.

### Brain structure (TBV) on socioeconomic status

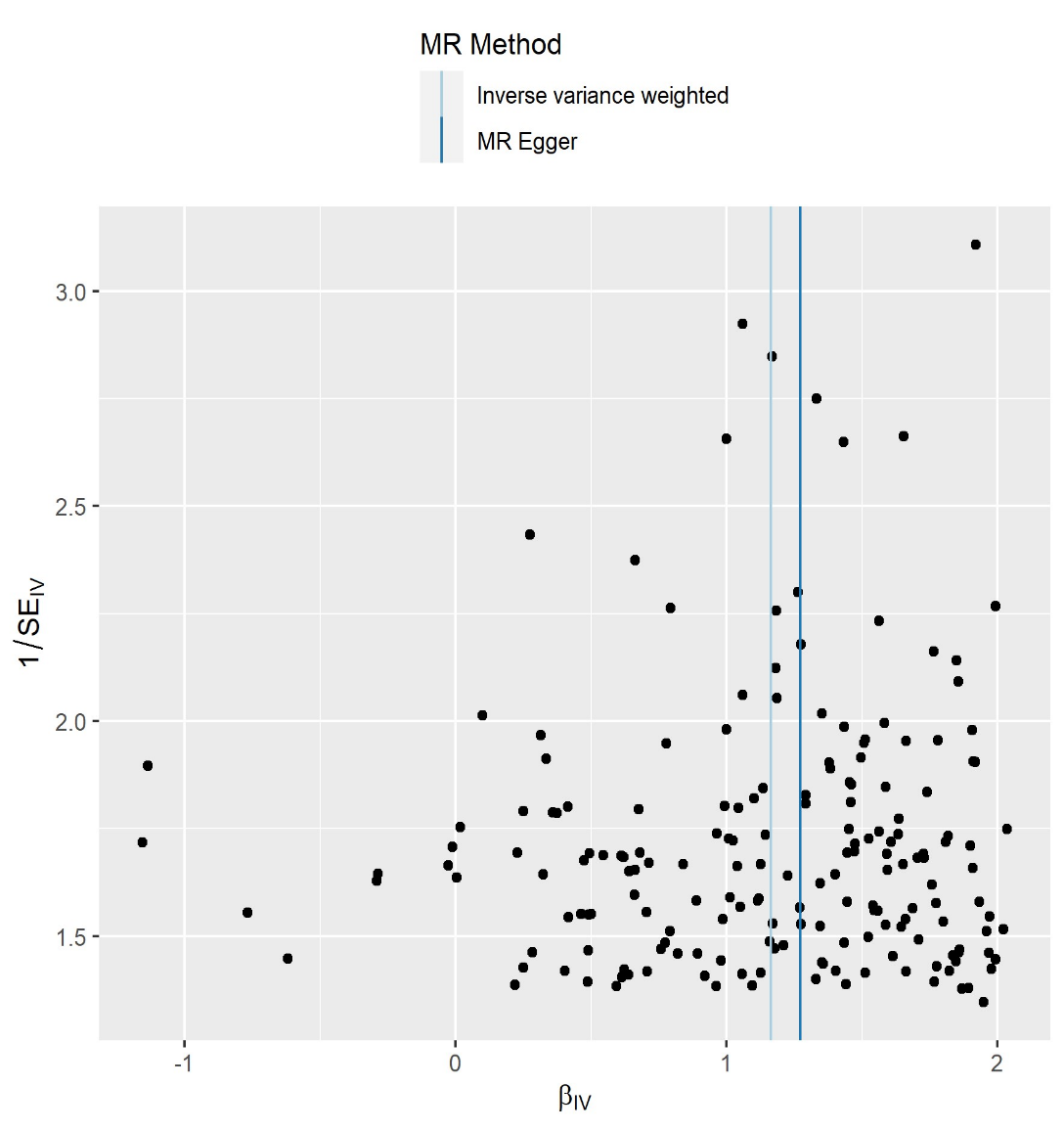

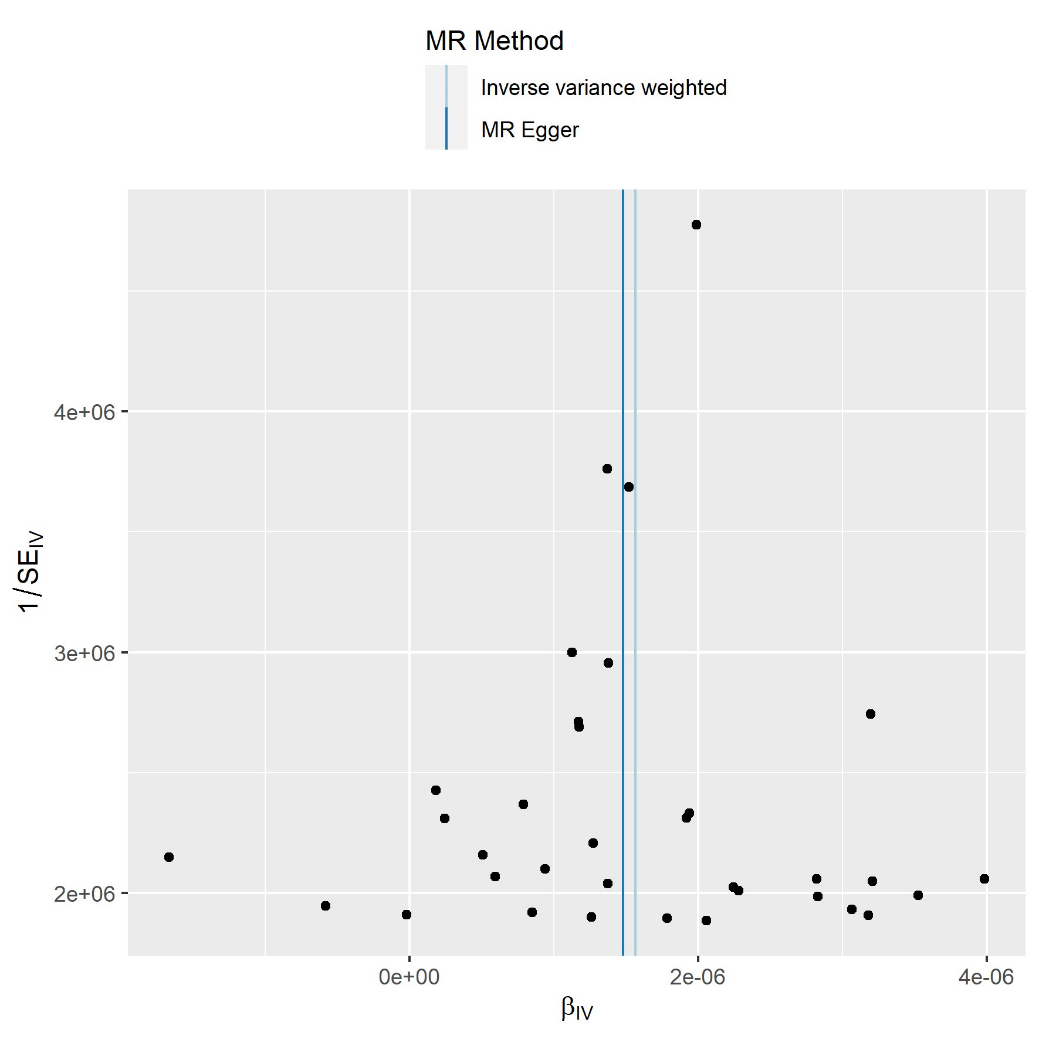

SES

**Supplementary Figure 3.** Funnel plot used to examine the extent to which pleiotropy is balanced across the instruments used in the univariate Mendelian randomisation of total brain volume on the general factor of socioeconomic status.

### Leave one out plot total brain volume on socioeconomic status

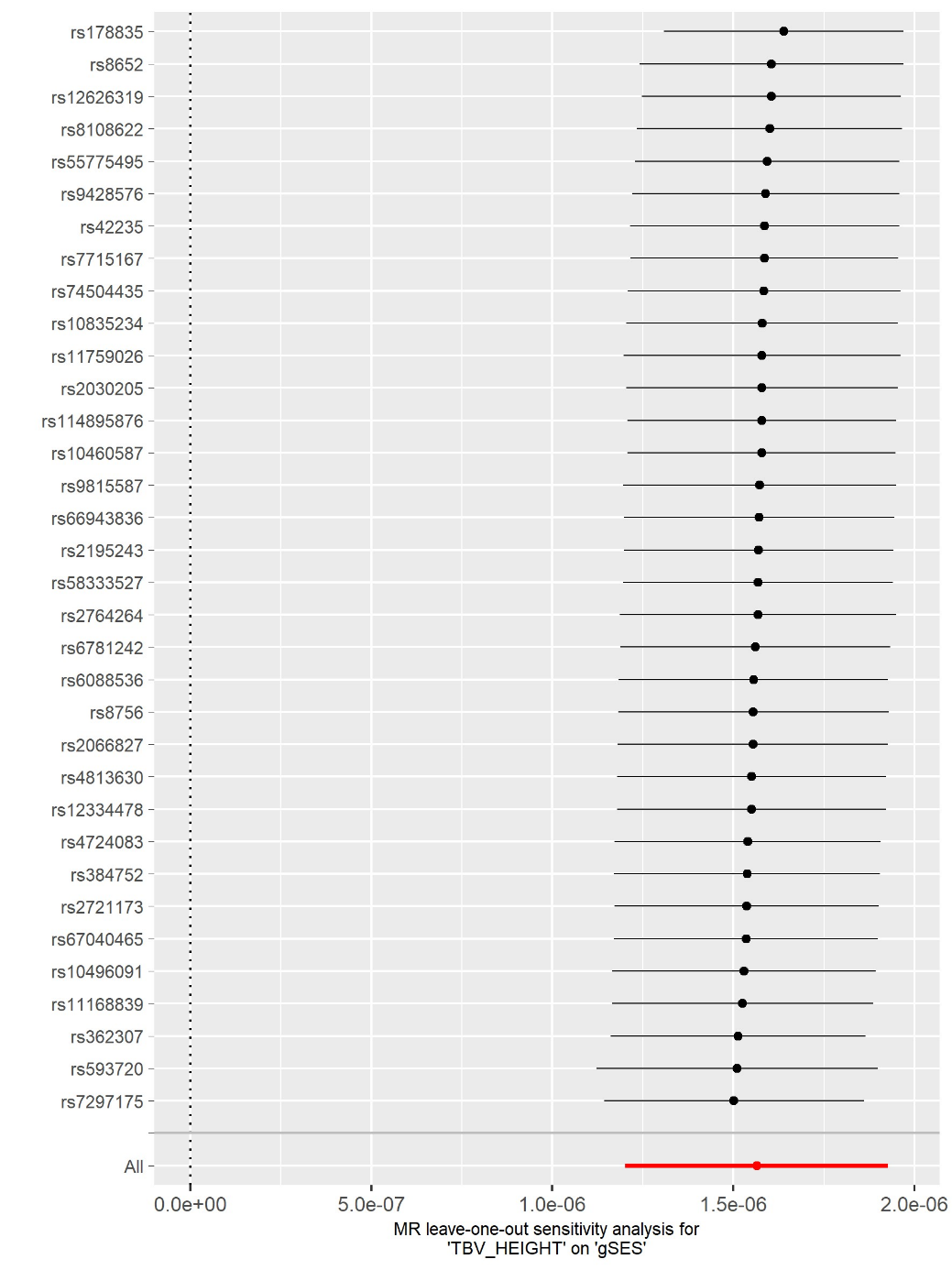

**Supplementary Figure 4.** Leave one out plot of the univariate Mendelian randomisation of total brain volume on general factor of socioeconomic status.

### Household income on brain structure

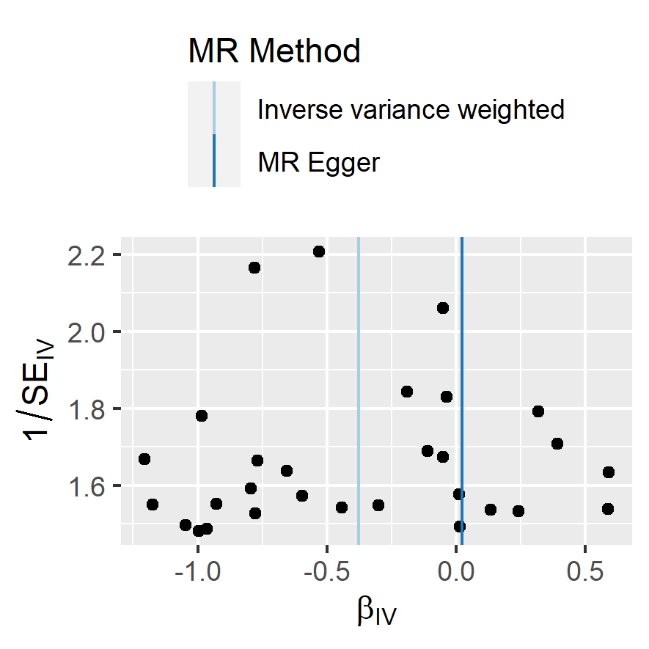

WMHicv

**Supplementary Figure 6.** Funnel plots used to examine the extent to which pleiotropy is balanced across the instruments used in the univariate Mendelian randomisation of Household income on brain structure.

### Educational attainment on brain structure

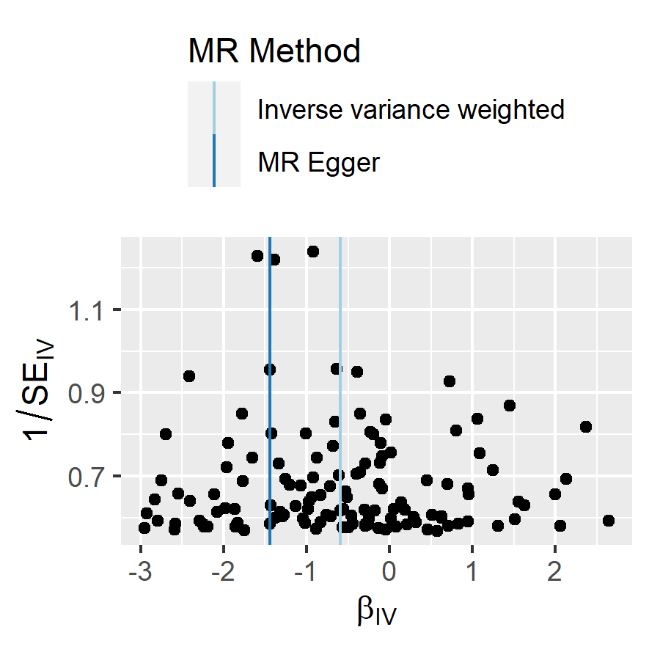

WMHicv

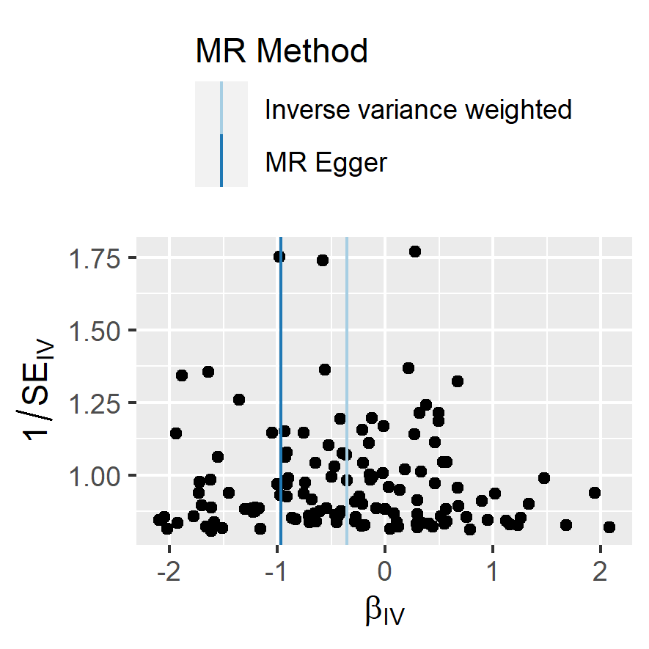

WMH

**Supplementary Figure 7.** Funnel plots used to examine the extent to which pleiotropy is balanced across the instruments used in the univariate Mendelian randomisation of Educational attainment on brain structure.

### Leave one out plots indicators of SES on Brain structure

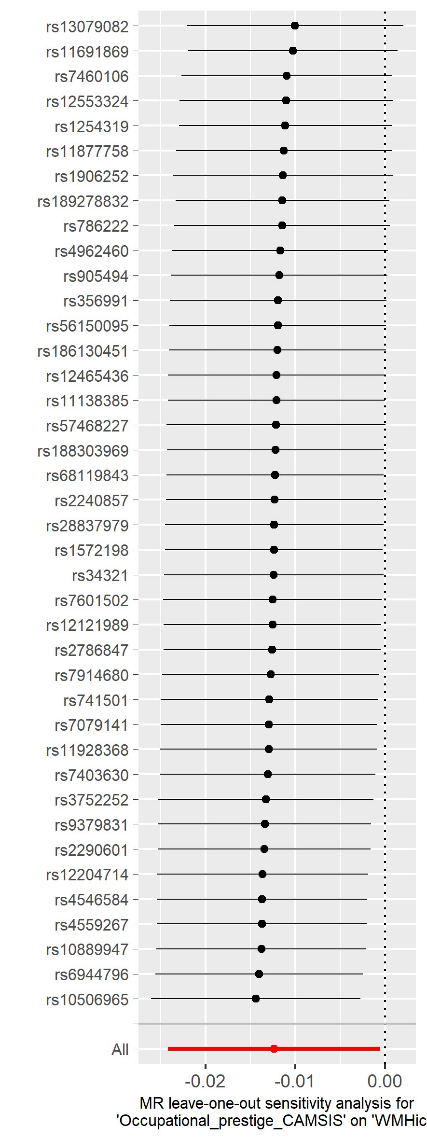

Occupational prestige

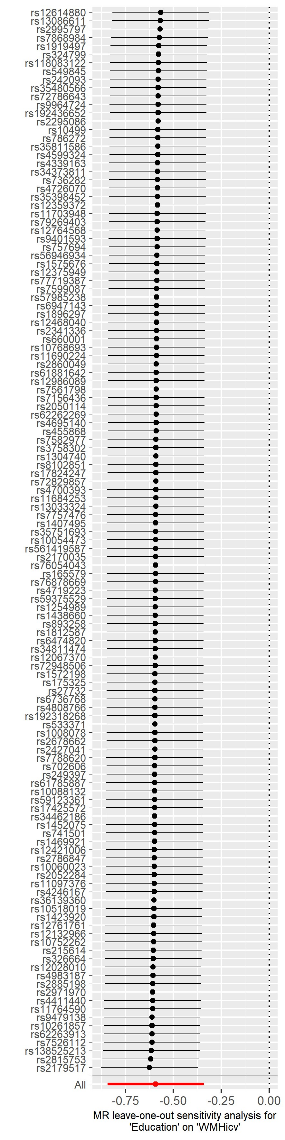

Educational attainment

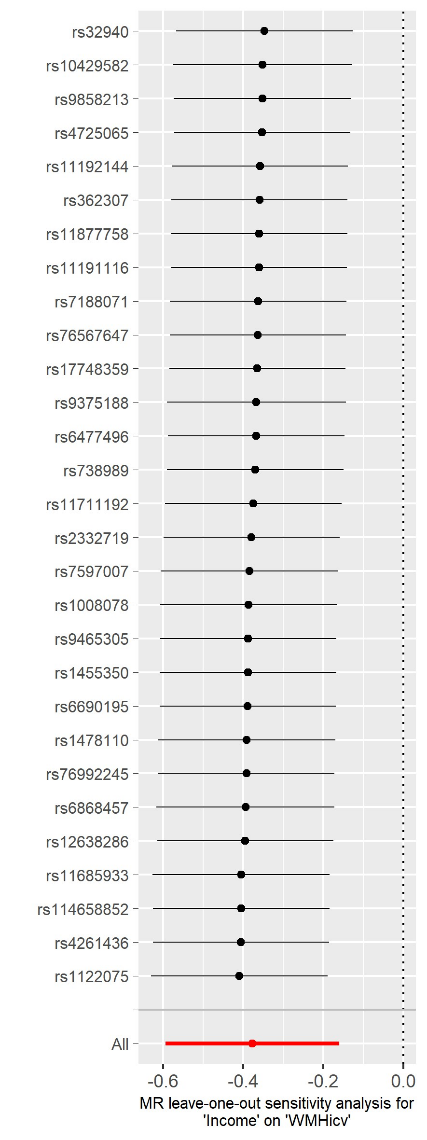

Household income

**Supplementary Figure 8.** Leave one out plot of the univariate Mendelian randomisation of Occupational prestige, household income, and educational attainment on white matter hyperintensities as a proportion of intracranial volume.

### Occupational prestige on brain structure

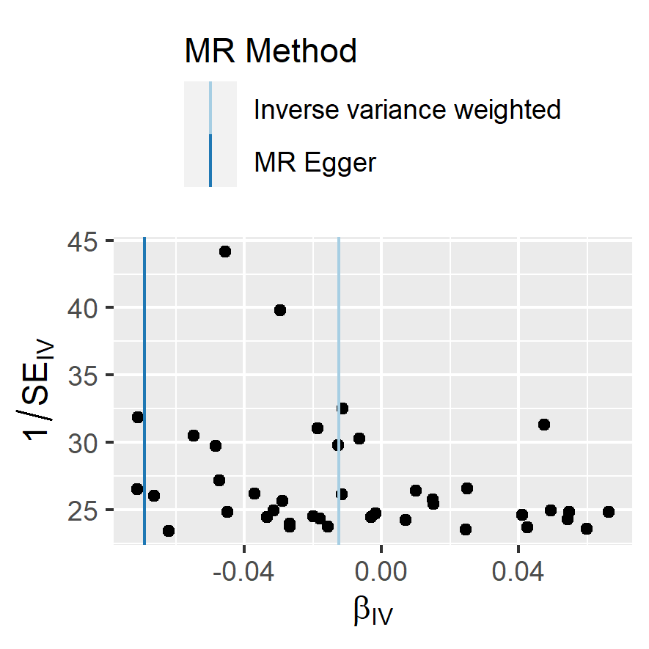

WMHicv

**Supplementary Figure 9.** Funnel plots used to examine the extent to which pleiotropy is balanced across the instruments used in the univariate Mendelian randomisation of Occupational prestige on brain structure.

### Social deprivation on brain structure

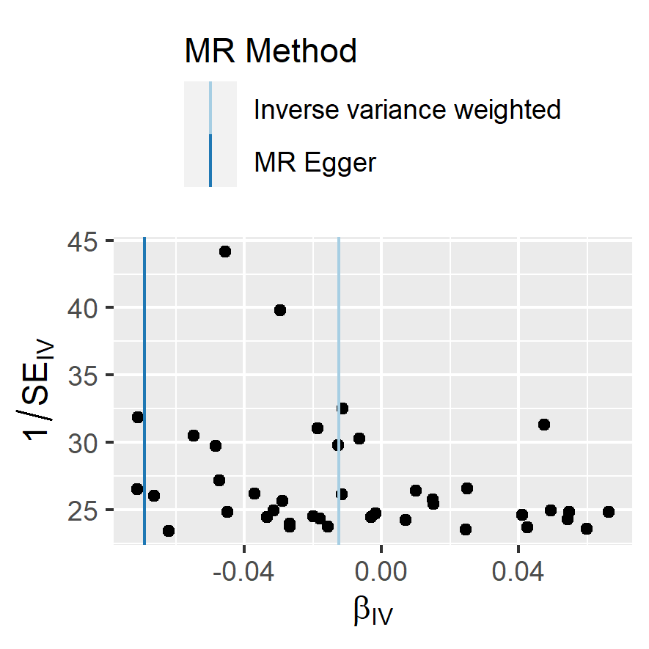

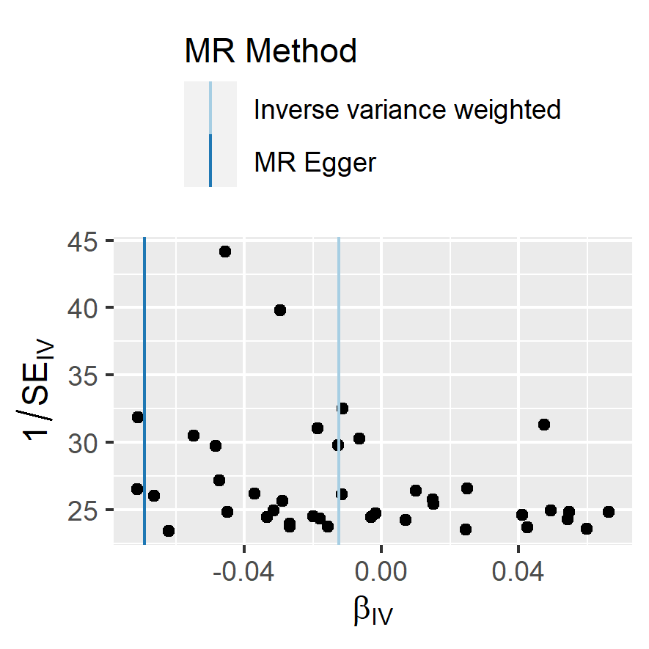

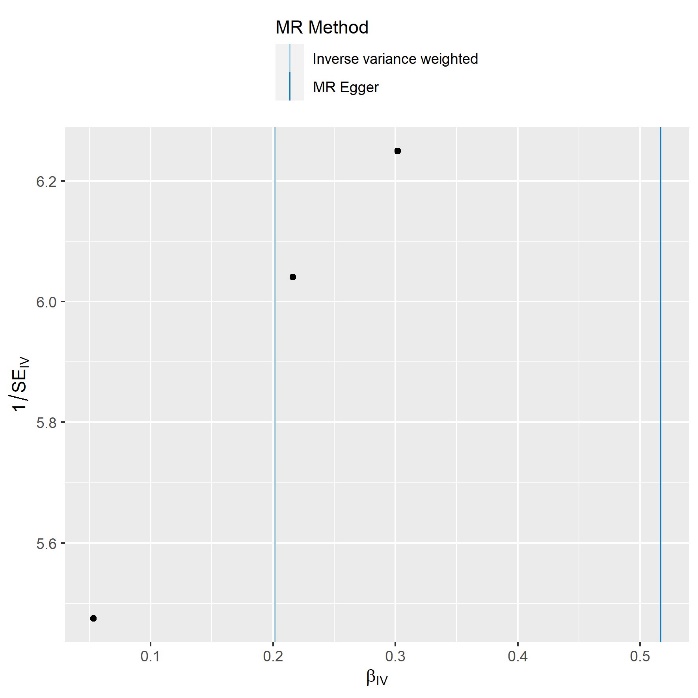

gMD

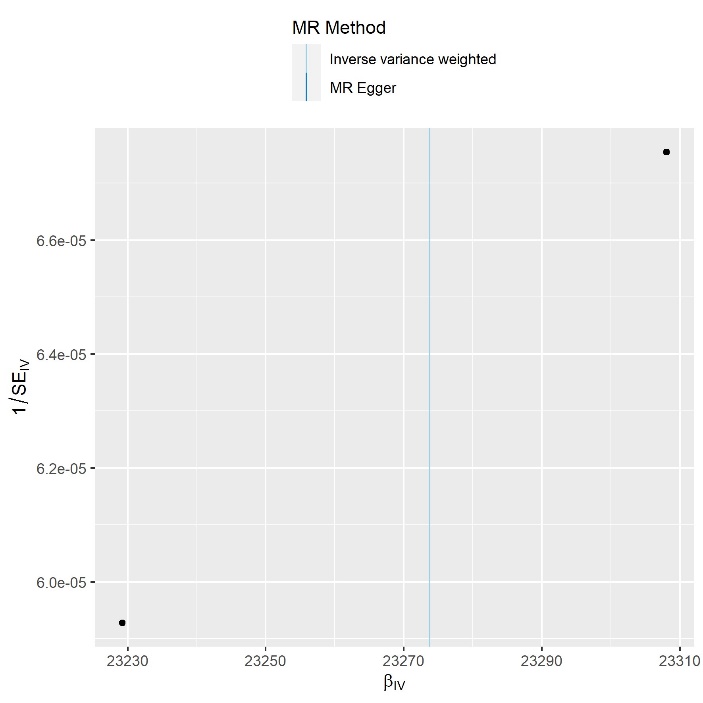

TBV

**Supplementary Figure 10.** Funnel plots used to examine the extent to which pleiotropy is balanced across the instruments used in the univariate Mendelian randomisation of Social deprivation on brain structure.

### Leave one out plot of Social deprivation on Brain structure

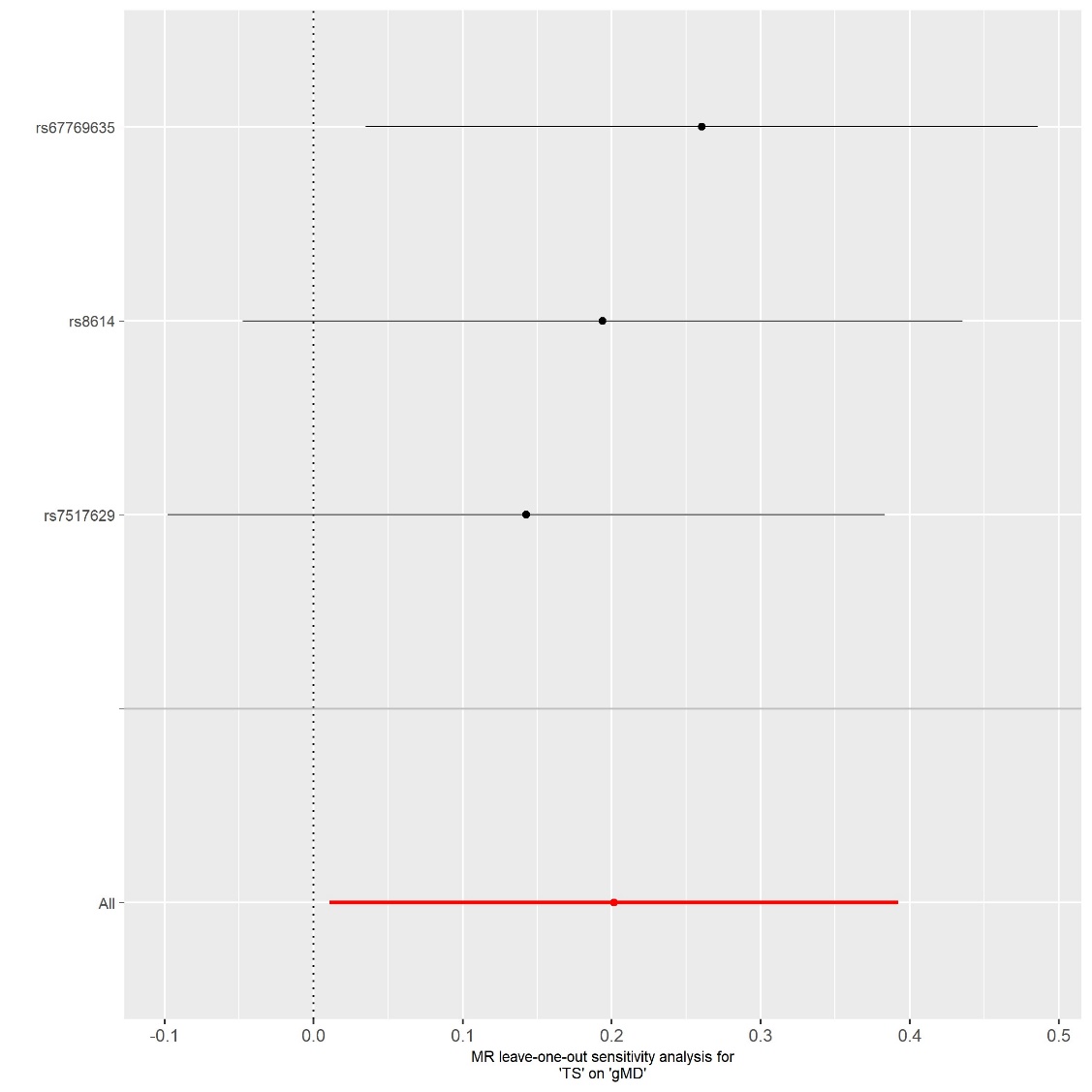

**Supplementary Figure 11.** Leave one out plot of the univariate Mendelian randomisation of Social deprivation on gMD.

### Total brain volume on indicators of socioeconomic status.

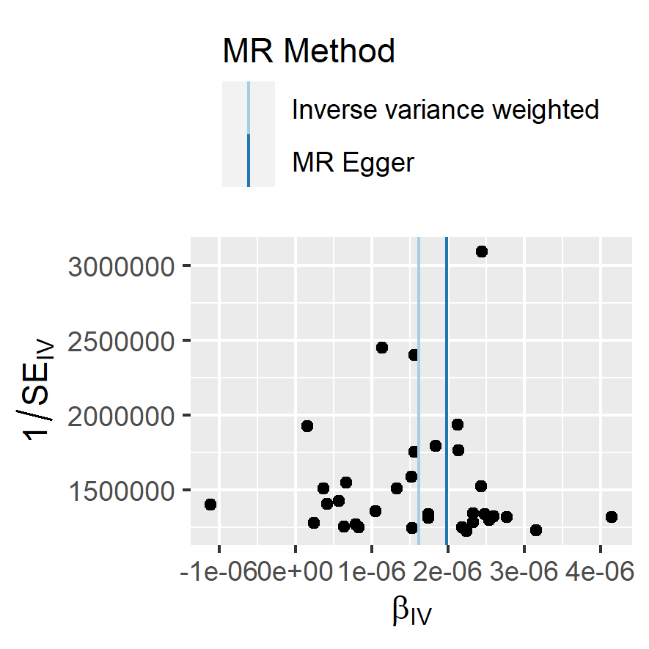

Income

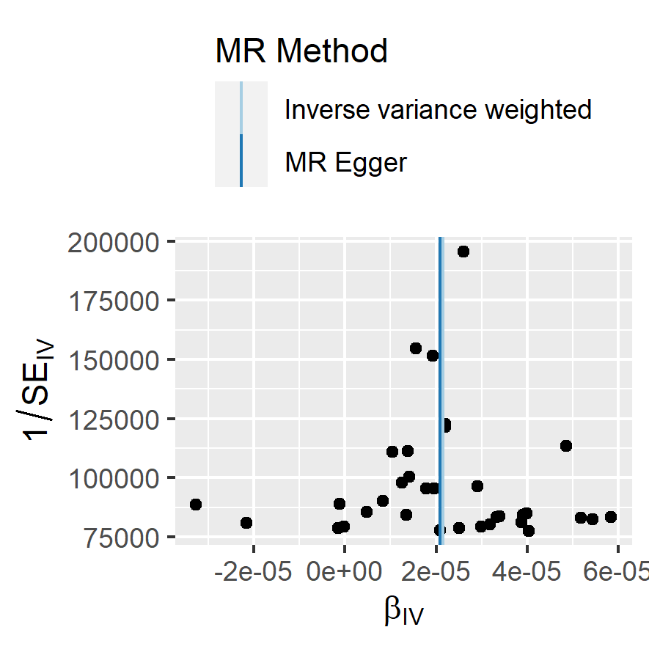

Occupation

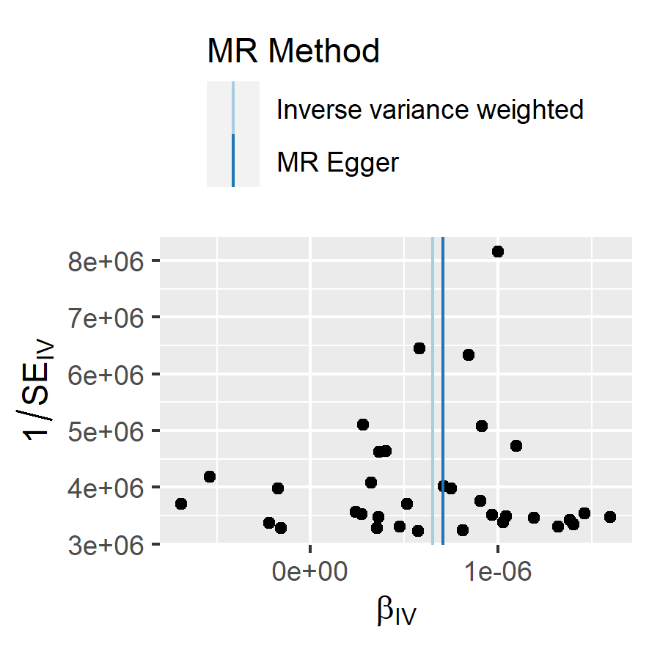

Education

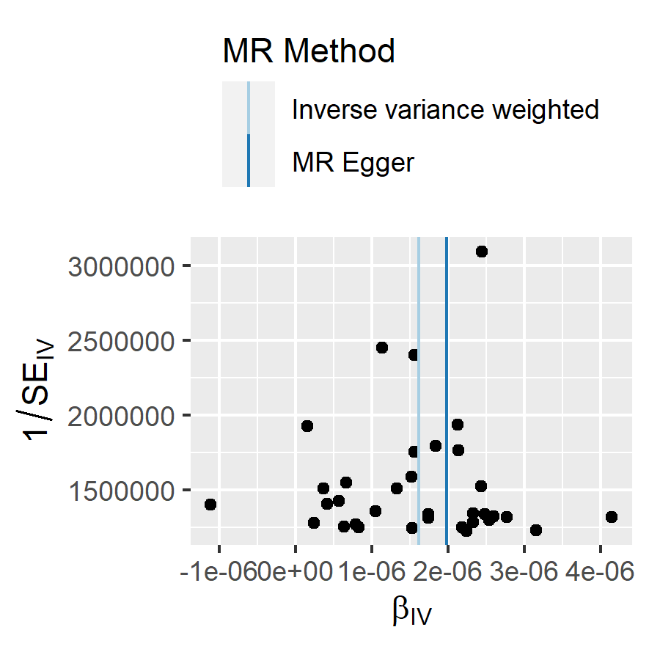

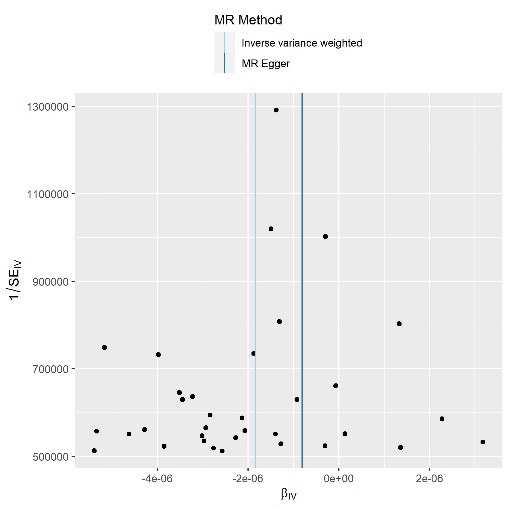

**Supplementary Figure 12.** Funnel plots used to examine the extent to which pleiotropy is balanced across the instruments used in the univariate Mendelian randomisation of total brain volume on indicators of SES.

### Leave one out plots indicators of total brain volume on indicators of SES

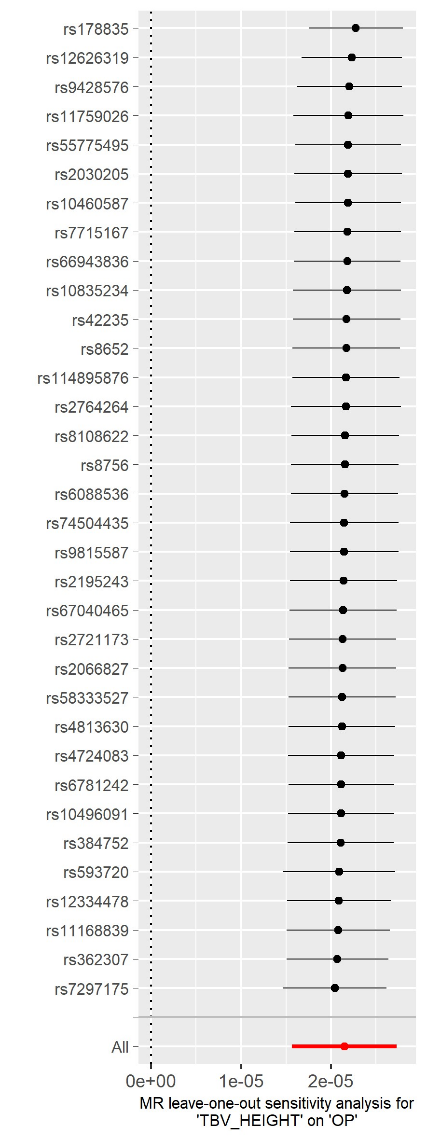

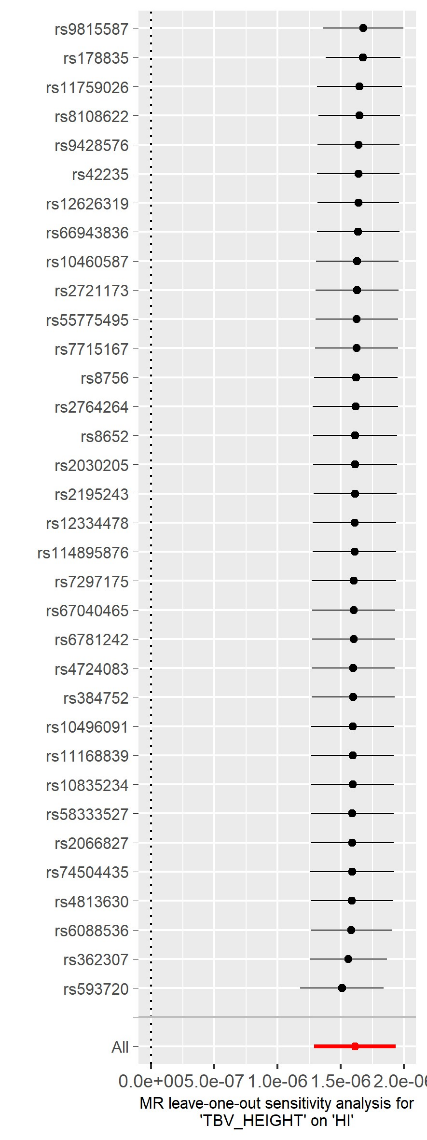

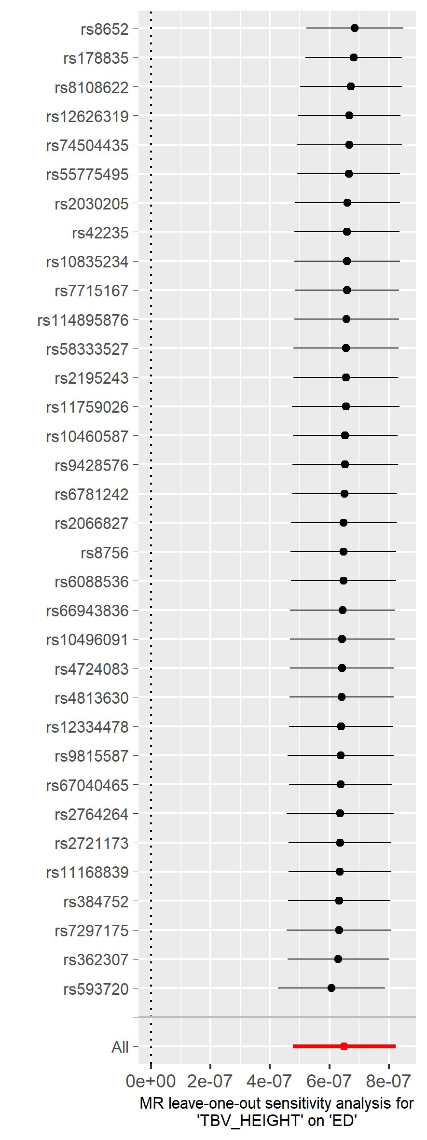

**Supplementary Figure 13.** Leave one out plot of the univariate Mendelian randomisation of total brain volume on occupational prestige, household income, and educational attainment.

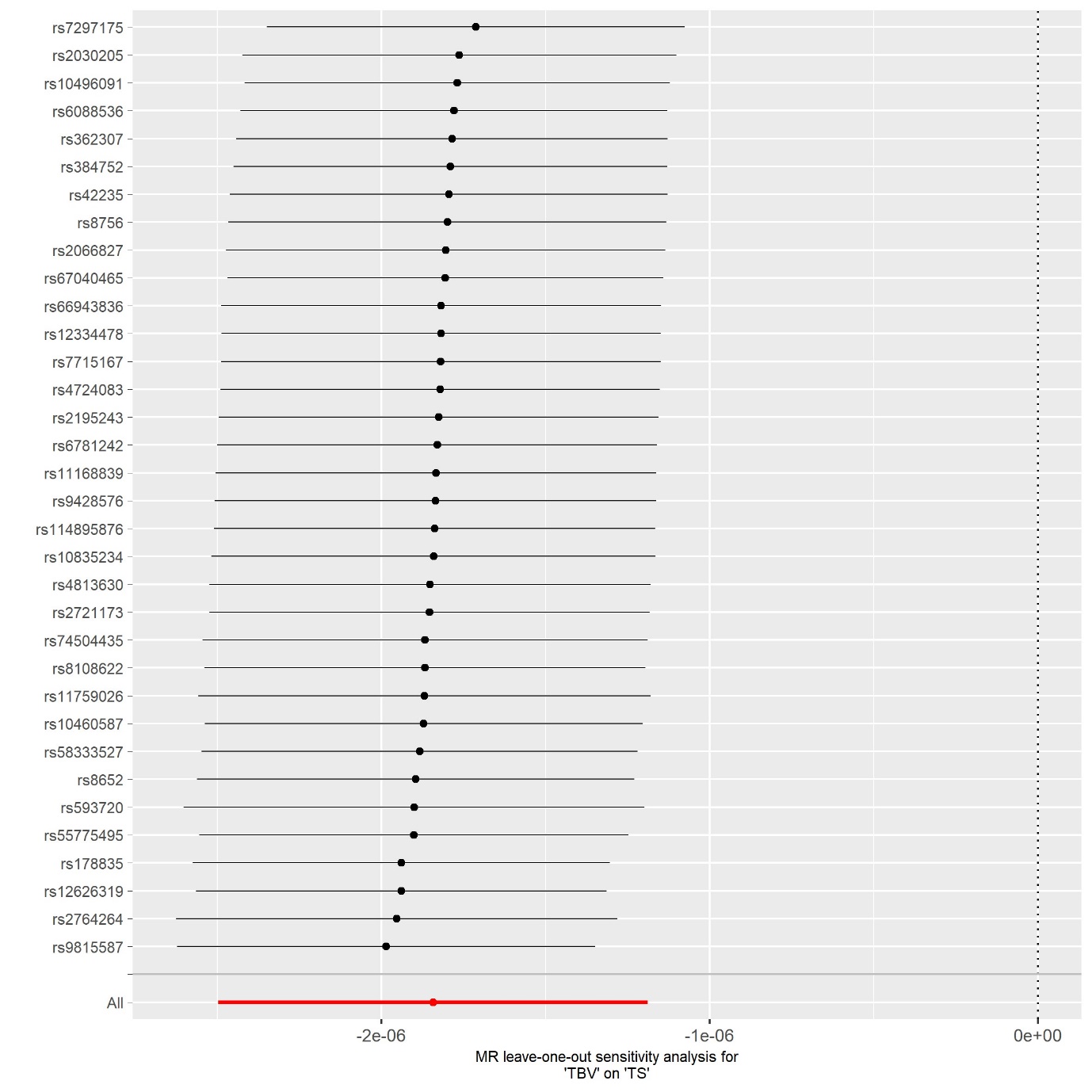

**Supplementary Figure 14.** Leave one out plot of the univariate Mendelian randomisation of total brain volume on social deprivation.

### Genetic overlap between intelligence and SES traits

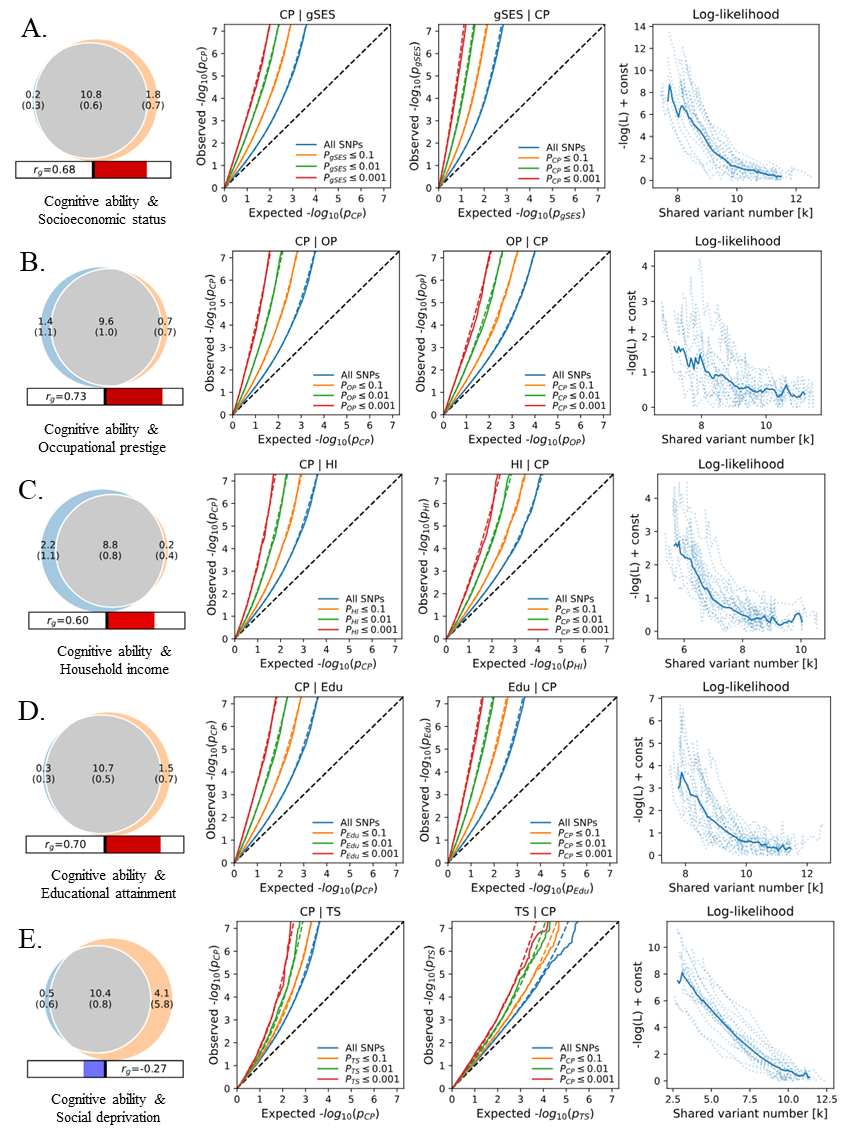

**Supplementary Figure 15.** Mixer results show the genetic overlap of cognitive ability (CP) with Socioeconomic status (gSES) in panel a, occupational prestige (OP) in panel b, household income (HI) in panel c, educational attainment (Edu) in panel d, and social deprivation (TS) in panel e. In each panel, the first plot shows the genetic overlap. Number of unique CP variants is shown in blue, number of unique SES variants is shown in orange, and number of shared variants is shown in grey. The unit is in thousand and the standard error of the estimate is in brackets. The bar beneath the Venn diagram shows the genetic correlation between CP and SES. The second plot is the Q-Q plot of CP GWAS condition on SES variants. The third plot is the Q-Q plot of SES GWAS condition on CP variants. The forth plot shows the negative log likelihood ratio of the genetic overlap model. Model with the lowest negative log likelihood ratio is defined as the best model.

### Cognitive ability on measures of socioeconomic status

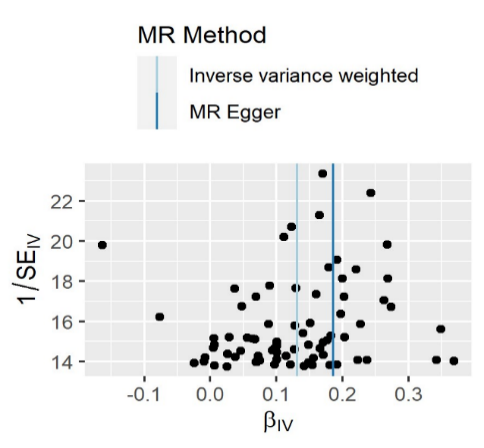

Socioeconomic status

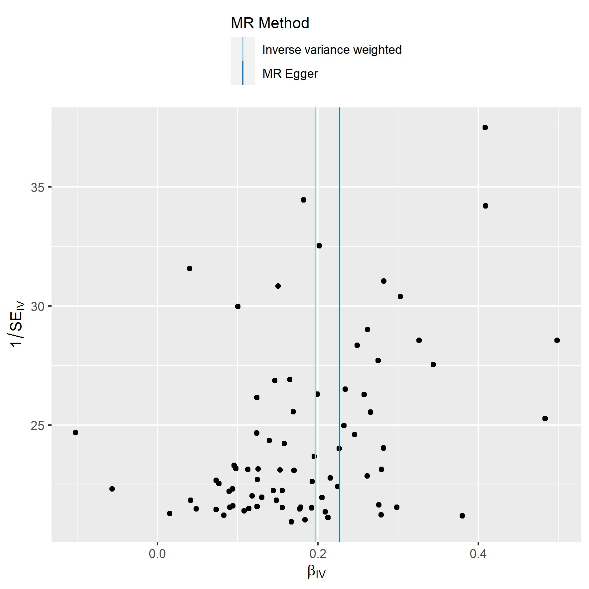

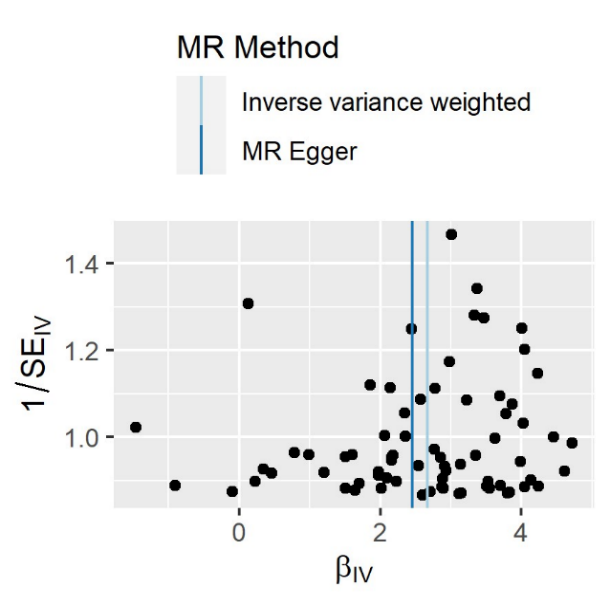

Occupational prestige

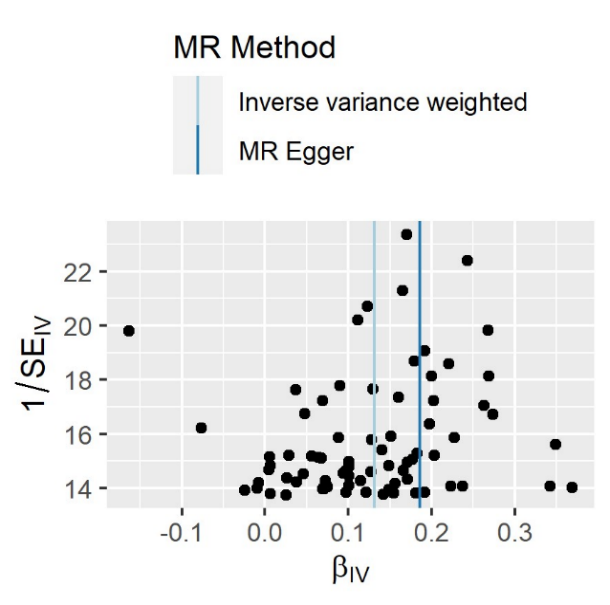

Household income

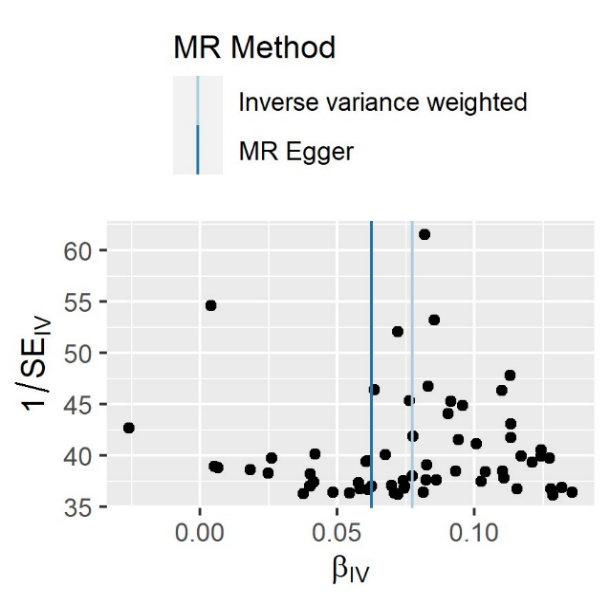

Educational attainment

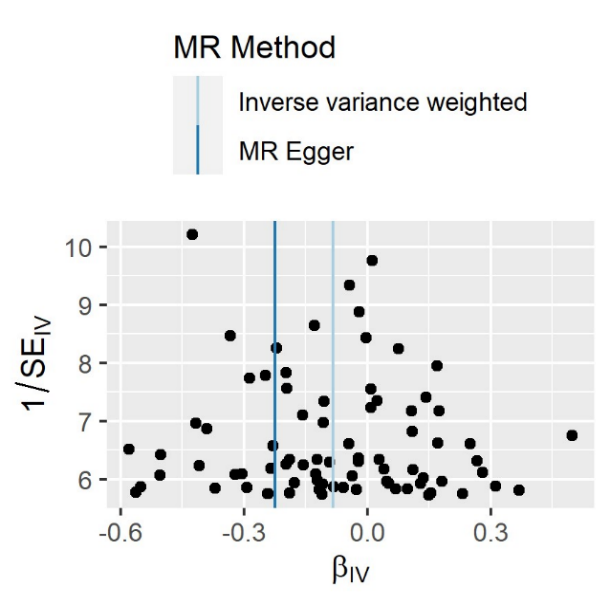

Social deprivation

**Supplementary Figure 16.** Funnel plots used to examine the extent to which pleiotropy is balanced across the instruments used in the univariate Mendelian randomisation of cognitive ability on the general factor of SES and on indicators of SES.

### Cognitive ability on measures of SES

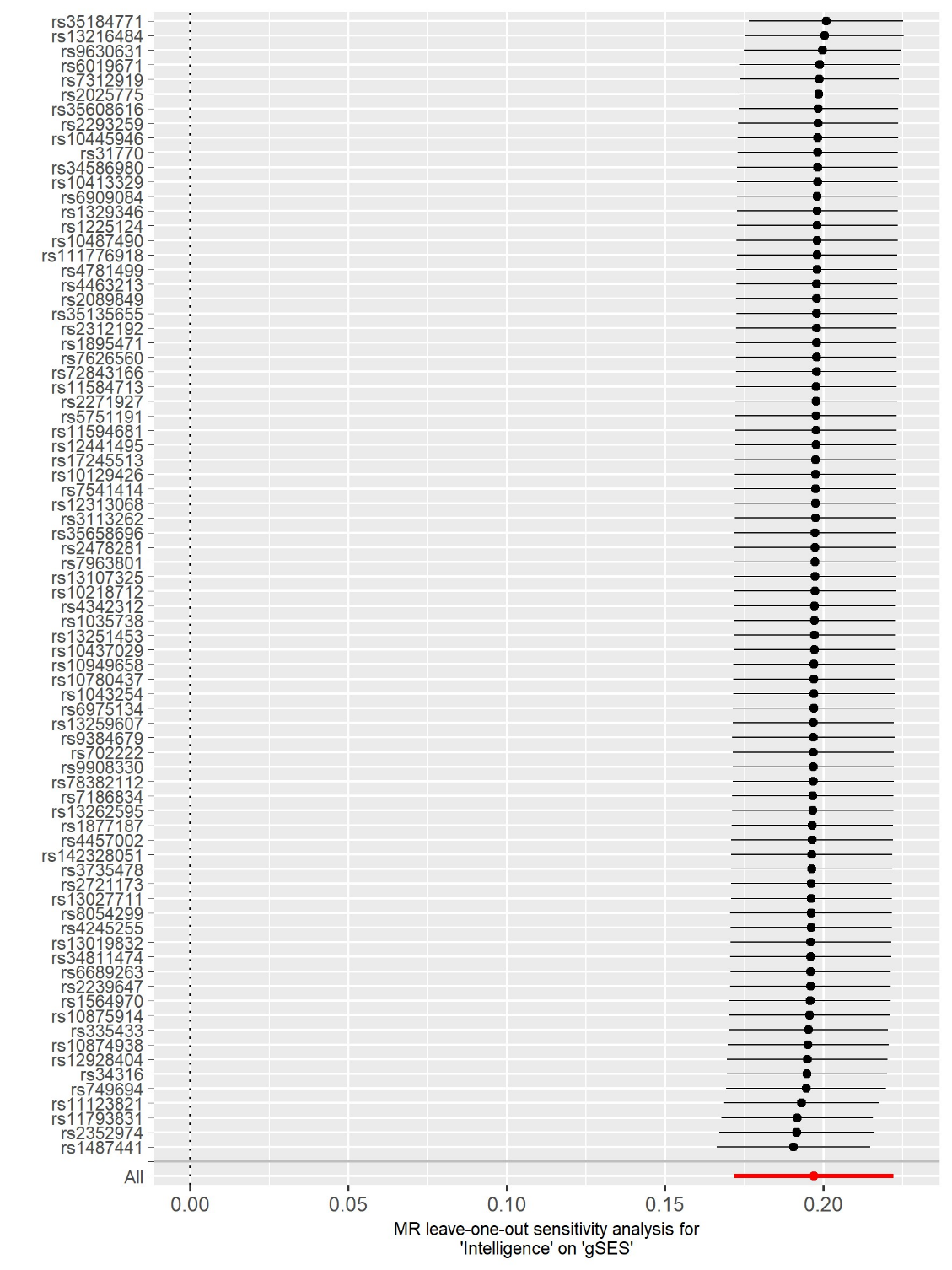

General factor of Socioeconomic status

**Supplementary Figure 17.** Funnel plots used to examine the extent to which pleiotropy is balanced across the instruments used in the univariate Mendelian randomisation of cognitive ability on the general factor of SES and on indicators of SES.

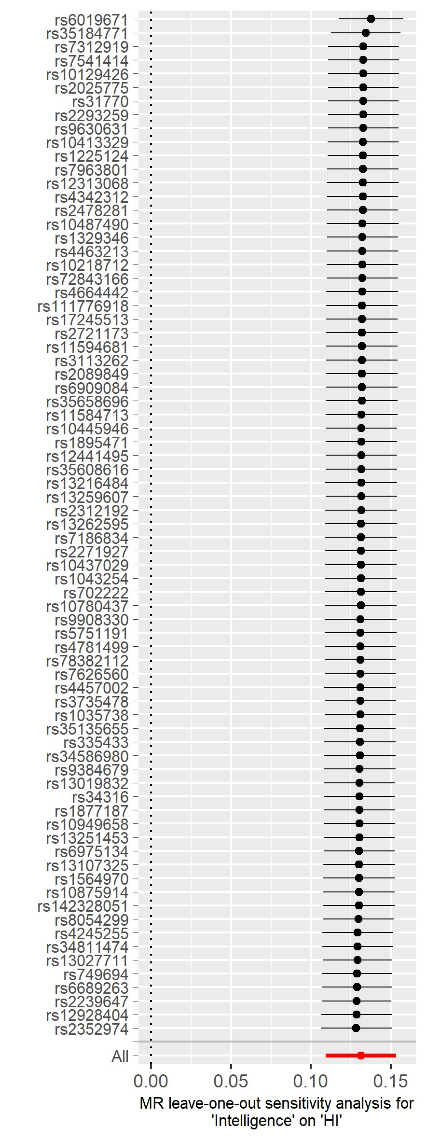

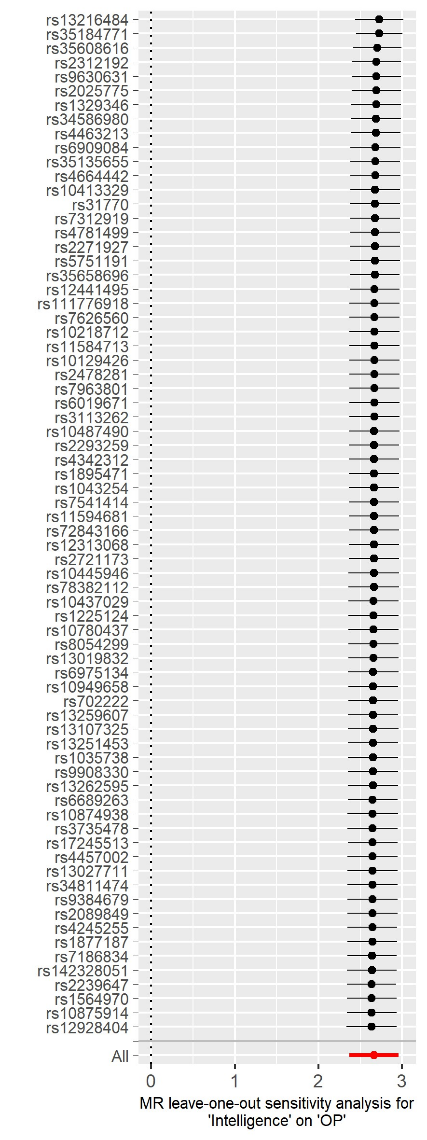

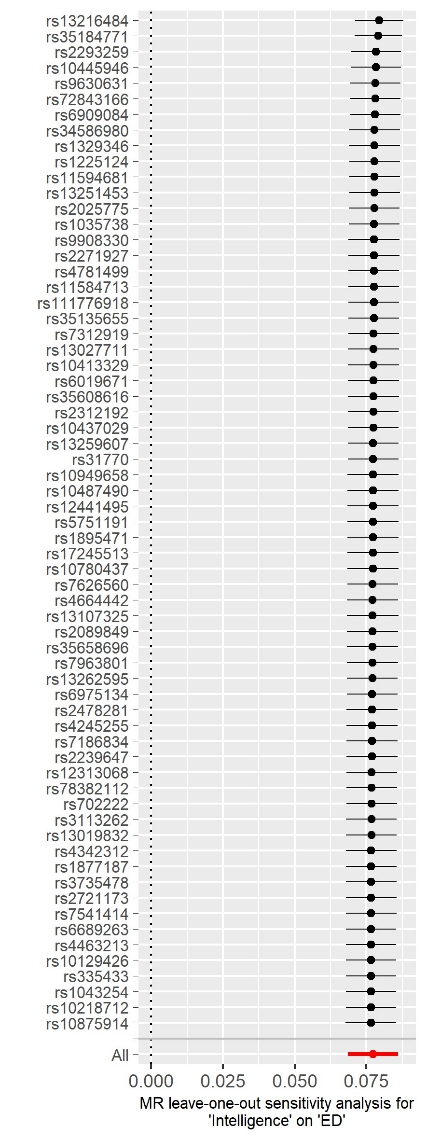

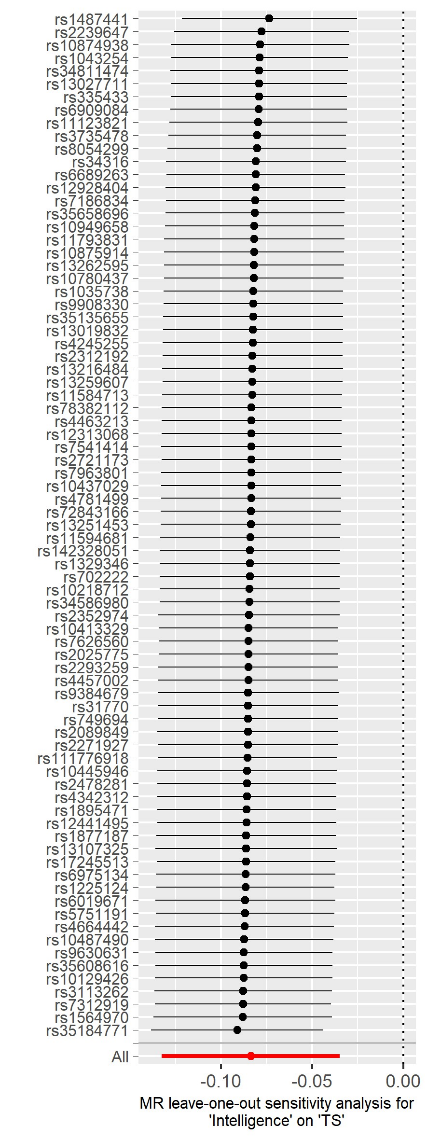

Occupational prestige

Household income

Educational attainment

Social deprivation

**Supplementary Figure 18**. Leave one out plot of the univariate Mendelian randomisation of cognitive ability on measures of SES.

### Socioeconomic status on Cognitive ability

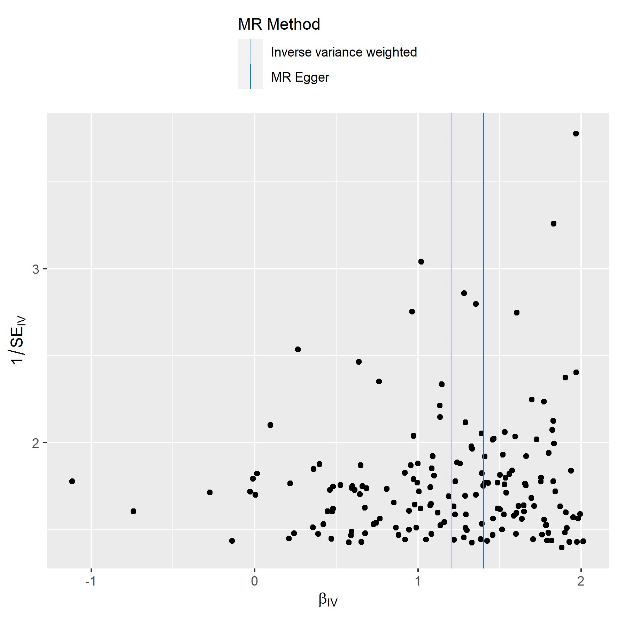

Socioeconomic status

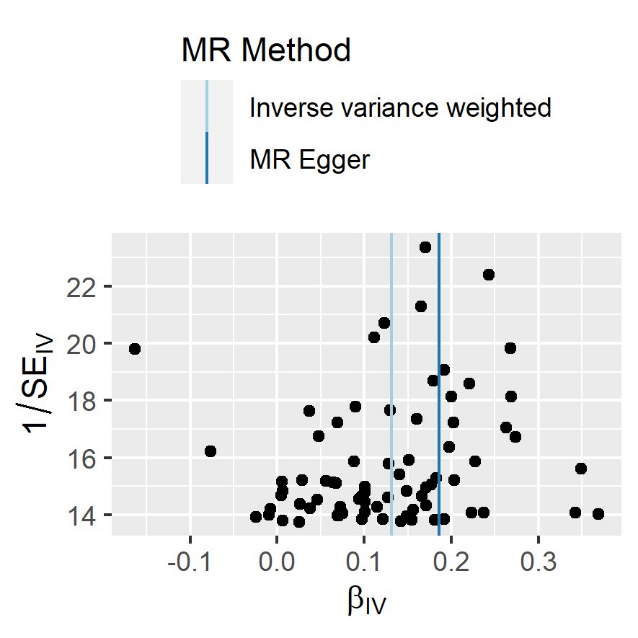

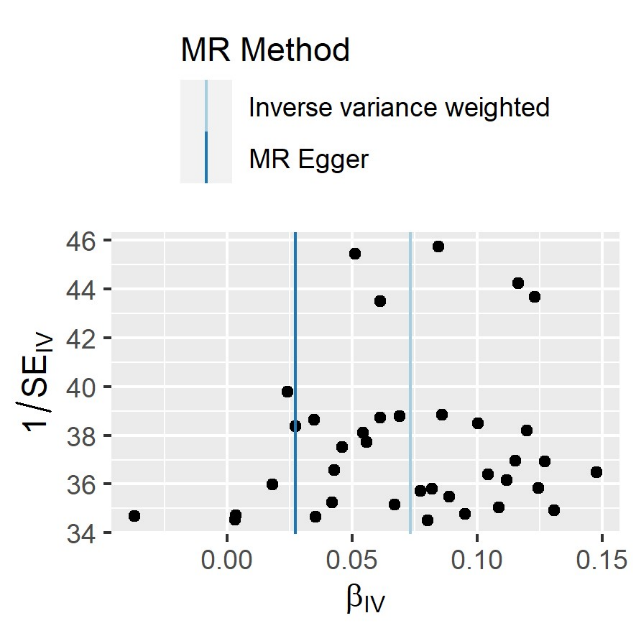

Occupational prestige

Social deprivation

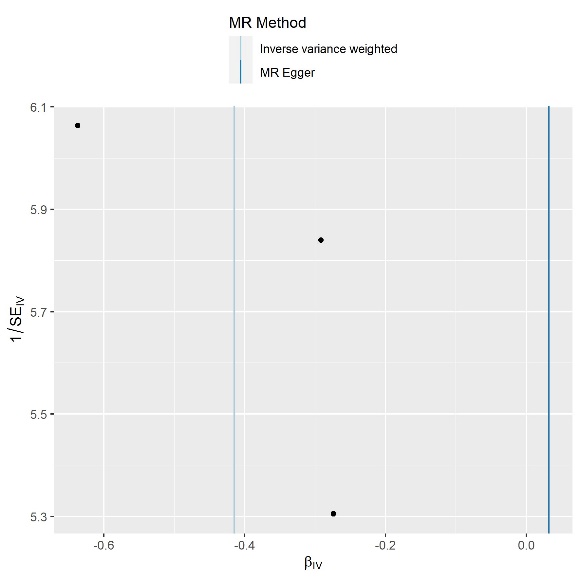

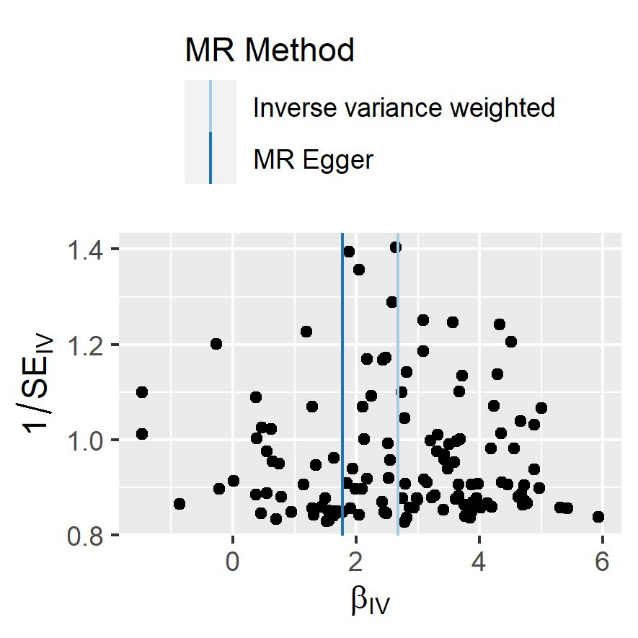

Educational attainment

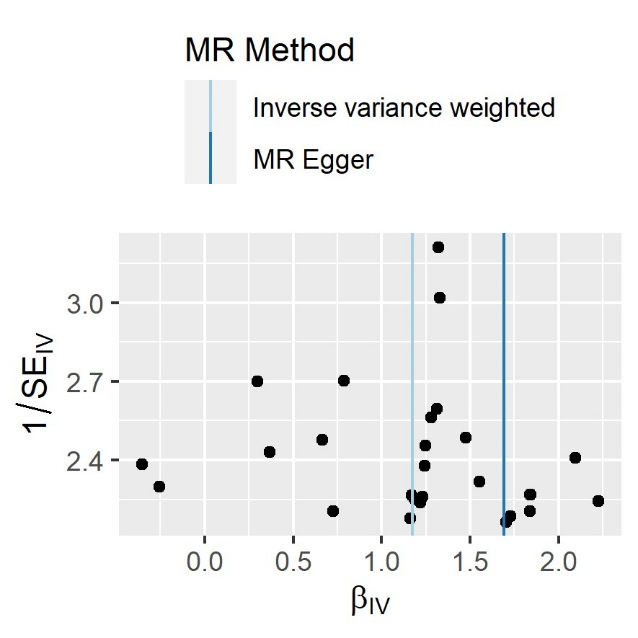

Household income

**Supplementary Figure 19.** Funnel plots used to examine the extent to which pleiotropy is balanced across the instruments used in the univariate Mendelian randomisation of the general factor of SES and indicators of SES on cognitive ability.

### General factor of SES on cognitive ability

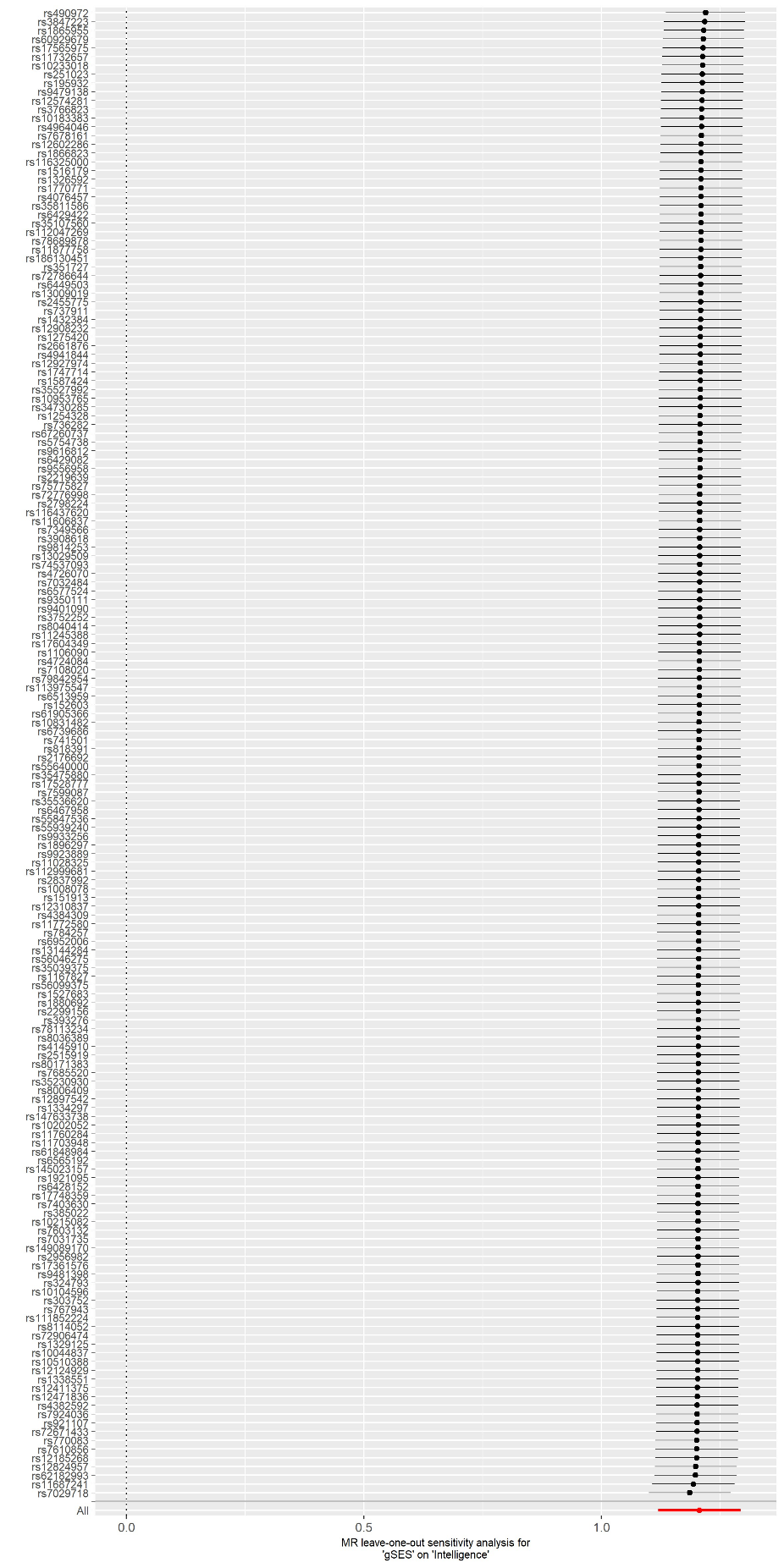

**Supplementary Figure 20.** Leave one out plot of the univariate Mendelian randomisation of the general factor of socioeconomic status on cognitive ability.

### Indicators of SES on cognitive ability

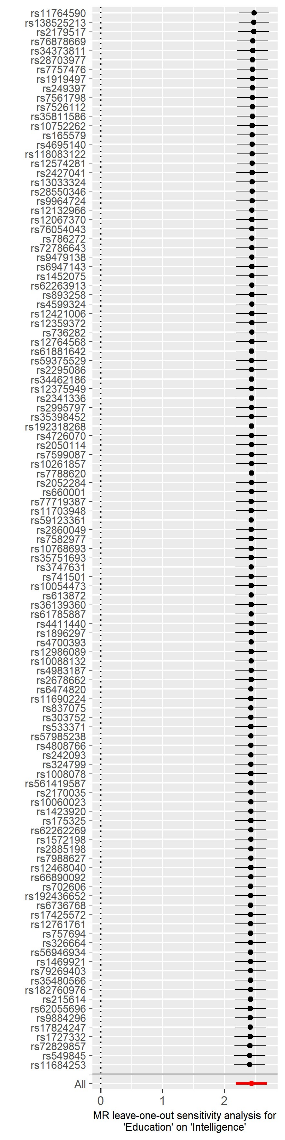

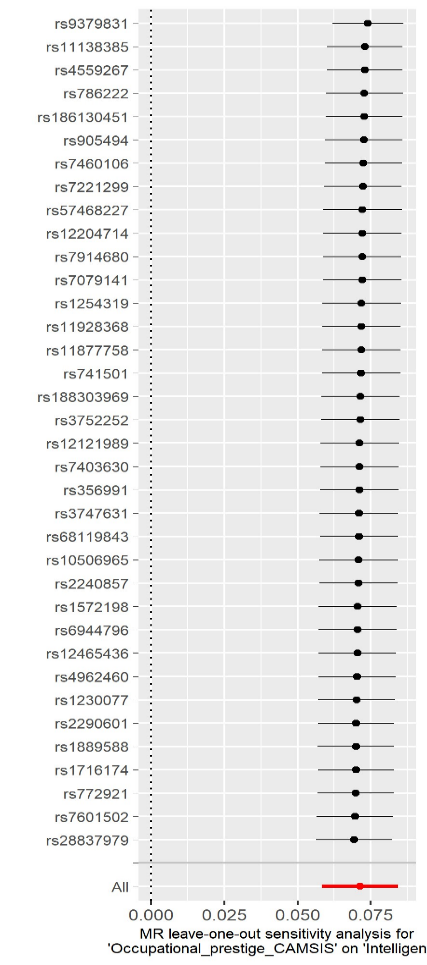

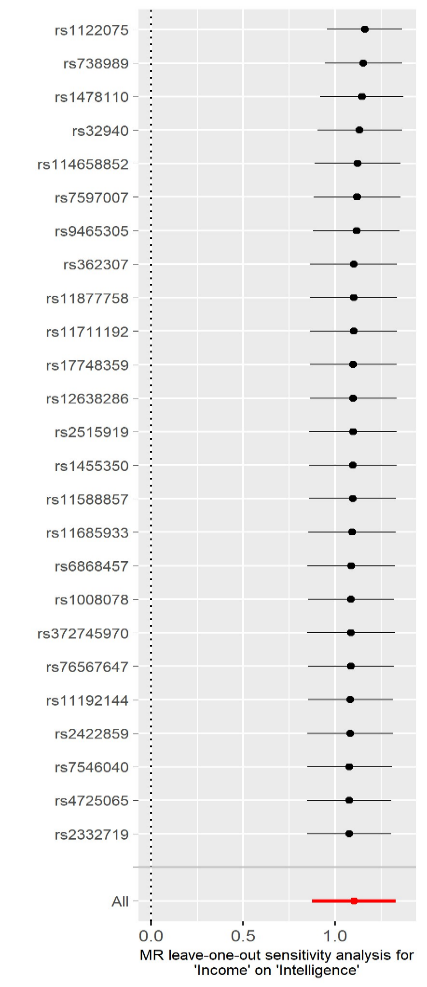

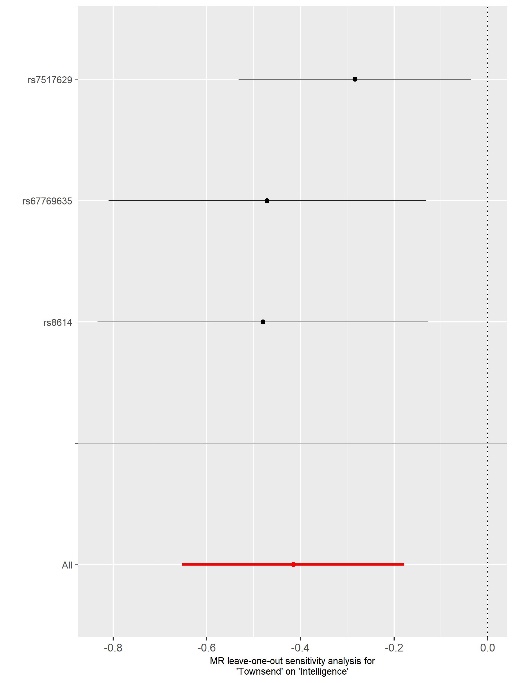

**Supplementary Figure 21.** Leave one out plot of the univariate Mendelian randomisation of the indicators of socioeconomic status on cognitive ability.

### Cognitive ability on brain structure

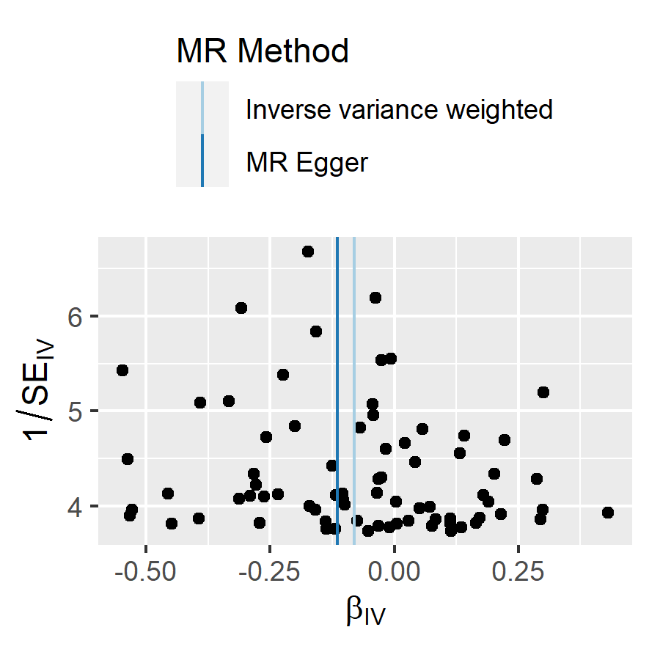

WMHicv

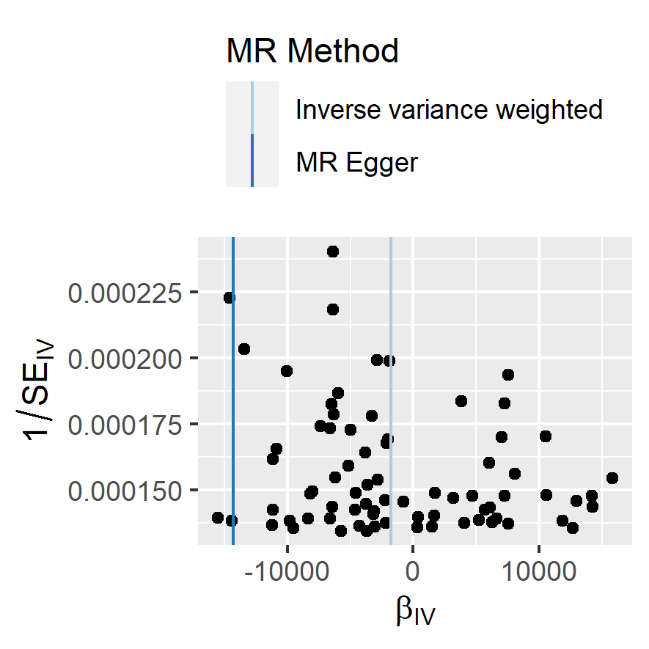

WMicv

**Supplementary Figure 22**. Funnel plots used to examine the extent to which pleiotropy is balanced across the instruments used in the univariate Mendelian randomisation of intelligence on brain structure.

### Total brain volume on cognitive ability.

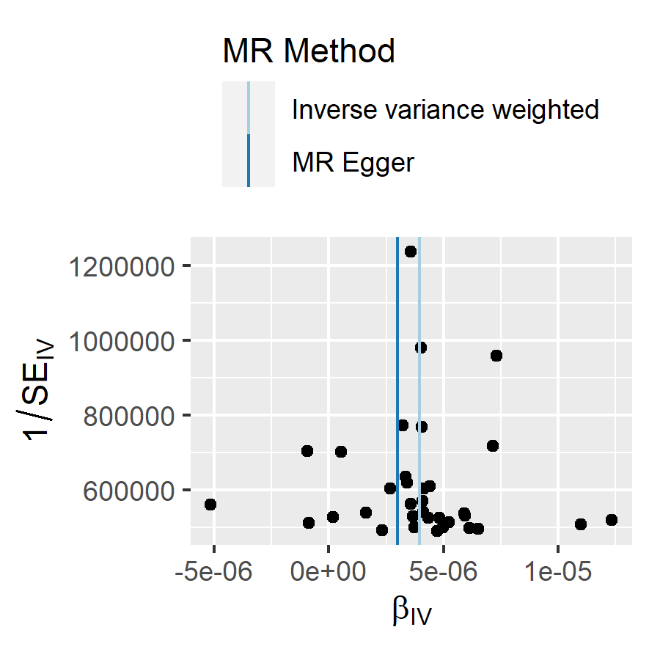

Intelligence

**Supplementary Figure 23.** Funnel plots used to examine the extent to which pleiotropy is balanced across the instruments used in the univariate Mendelian randomisation of total brain volume on cognitive ability.

### Bidirectional effect of brain structure on cognitive ability

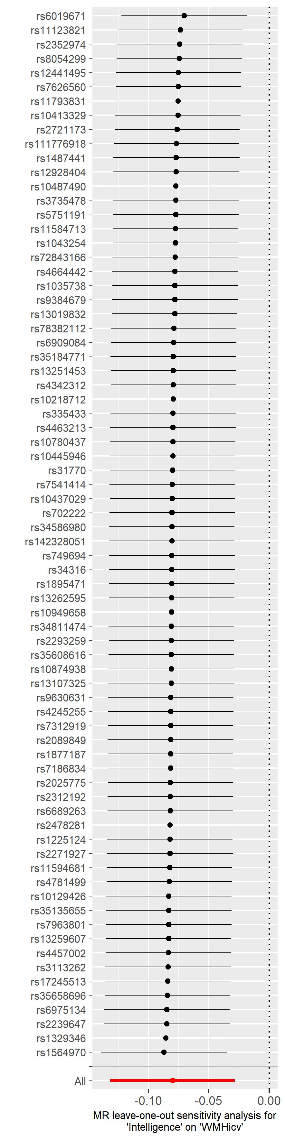

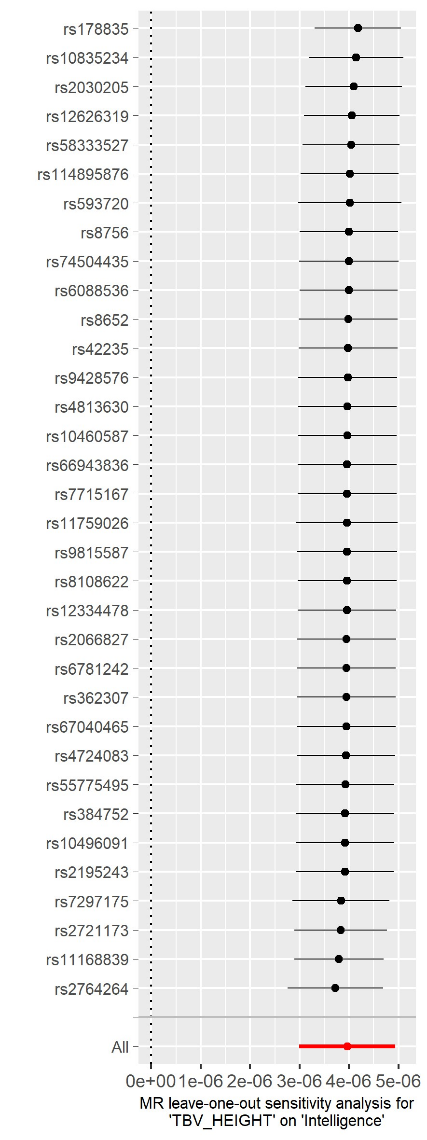

**Supplementary Figure 24**. Leave one out plots of the univariate Mendelian randomisation of cognitive ability on WMHicv, and total brain volume on cognitive ability.
