## Supplementary Notes for "Deciphering the Influence of Socioeconomic Status on Brain Structure: Insights from Mendelian Randomization"

Frequently Asked Questions (FAQs) regarding:

### What was the goal of this study?

Our goal was to examine the causal effects of socioeconomic status (SES) on brain structure and function. In addition, as differences in SES are also associated with differences in cognitive ability, we investigated if these differences in cognitive ability accounted for the effects we found of SES on brain structure.

### Why look at the effect of SES on brain structure?

Our investigation into the links between social inequality and neuroscience will facilitate our understanding of the brain in a number of substantive ways. First, our work has the potential to identify mechanisms through which brain structure and function relates to differences in social inequality (as measured by income, occupation, education, social deprivation). This would complement other areas of research linking differences in SES to health behaviours, access to economic resources and medical care.

Second, our efforts in understanding how differences in SES relate to brain structure and function could help to improve replication by providing explanations into replication failures. For example, the association between brain size and cognitive ability is well known but this association varies across countries. Such findings may indicate that differences in social factors are, in part, driving the link between brain volume and cognitive ability.

### How does one look for causality?

One method used by scientists to examine causality is through a randomised control trial (RCT). An RCT can be used to gauge whether a particular treatment or medical intervention causes a reduction in disease or disease prevalence. In an RCT, participants will be randomly divided into at least two groups one of which would receive the treatment or intervention whereas the other would not. As participants are allocated to groups at random, any other factors that effect disease outcome should be equally present in both groups. Thus, in an RCT any difference in disease prevalence between the two groups can be attributable to the causal effect of the treatment/intervention.

However, there are instances of where an RCT can not be conducted. For example, when examining the effect of SES, as individuals cannot be randomly assigned to an SES group as this would be deeply unethical.

### How did this study examine causality?

In our study used an instrumental variable (IV) to capture differences in SES. An IV is related to the exposure (in this instance SES) but not to the outcome (brain structure and function) except through its association with the exposure. Instrumental variables can be used to examine causality when they are randomly assigned to members of a population. In our study we made use of a naturally occurring event whereby genetic variants are seen to be randomly assigned at conception. These genetic variants can then be used as instrumental variables as they capture differences in SES. Using genetic variants as instrumental variables to examine the evidence of a causal relationship is called Mendelian randomisation (Figure 1.).

**Figure 1.** Showing the similarity between a randomised control trial (RCT) and a Mendelian randomisation (MR) study design. In an RCT participants are randomly assigned into two groups one of which will receive a treatment whereas the other will receive a placebo (for example). In an MR design genetic variants are seen to be randomly allocated at conception and so by using genetic variants as instrumental variables for SES differences and so SES can be randomised in a population.

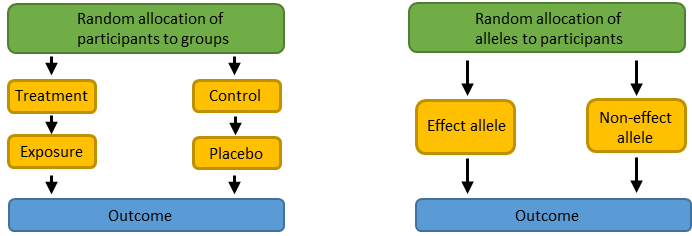

### How was socioeconomic status measured in the current study?

We used four indices of SES: occupational prestige, household income, educational attainment, and social deprivation. Occupational prestige was derived using what job participants have. Household income was measured as the income in the household of each participant prior to tax. Educational attainment was measured in two different, but related, ways. In one group, educational attainment was measured as a binary trait where those who attained a university level degree were coded as 1 and those without a university level degree were coded as 0. In our replication group educational attainment was measured as the number of years of education each participant attained. Social deprivation was measured using the Townsend Deprivation Index. This index uses data from the National census and information on unemployment, non-car ownership, non-house ownership, and overcrowding. In addition, we performed a multivariate analysis that allowed us to distil genetic effects into those that acted on each of these four indices of SES and those that were specific to anyone.

### How did you find genes associated with each indicator of SES?

We used a technique called a genome-wide association study (GWAS). A GWAS is a technique that examines if genetic variants are linked with a specific outcome. They have been extensively used to examine health conditions such as cardiovascular disease and major depressive disorder, as well as non-disease traits such as height, weight and cognitive ability. Our GWAS were conducted to examine four traits capturing different aspects of SES: occupational prestige, household income, educational attainment, and social deprivation.

The genetic variants tested in a GWAS are called single nucleotide polymorphisms (SNPs, pronounced “snips”). These are the most common source of genetic differences between people and describe which nucleotide (termed allele and describe which one of the four chemical letter an individual has: A, C, T, and G) a person has. SNPs usually are bi-allelic (i.e., binary). For example, in a population the “A” nucleotide may be present in 25%, with the remaining individuals having the “G” version. A GWAS study lines all the SNPs an individual has and tests the degree to which each one is linked to the outcome of interest. In the case of height, for example, each SNP may contribute only a tiny fraction towards height differences by contributing towards bone length, or hormone production.

In our study we performed four GWAS examining different indicators of SES: occupational prestige (n = 242,776), household income (n = 327,402), educational attainment (n = 377,477) and social deprivation (n = 382,030). We also used a statistical technique to determine if genetic effects were shared across these four measures of SES or if there were genetic effects that were unique to anyone of them.

### What did you find?

As expected, we found that the majority of the reasons why people differed in their level of SES was not due to genetic factors. We believe that these reasons are likely to be environmental in origin. What we mean by environmental would cover such things as social conditions, specific policies, and even luck. However, we did find genetic associations between genetic variation and differences in SES. We will speculate as to why these effects were found in the answer to our next question. First, we will describe the results in more details here.

For each indicator of SES we found evidence of a small genetic effect variation across all SNPs taken together accounted for 11% of occupational prestige, 7% of household income, 13% of educational attainment, and 4% of social deprivation. This is a small proportion of the total differences in SES and other methods that capture less common forms of genetic variation have been able to estimate that genetic effects account for around 40-50% of the variation in income and education respectively. Therefore, the estimates we present here are likely to be underestimates of the true extent that genetic differences are linked to differences in SES.

We found that the genetic effects for each of these four indicators of SES were largely the same and this allowed us to derive a general factor of SES. In total, genetic variation accounted for 11% of the differences in this general factor of SES. We next distilled these genetic effects into those that were shared across each of the four measures of SES and those that were common to any specific indicator (occupational prestige, household income, occupational prestige, or social deprivation). We found that of the 399 regions of the genome (termed loci) that were associated with this general factor of SES only two showed evidence that their association was being driven by a subset of the four indicators rather than capturing what was common across all four. This is strong evidence that the genetic factors associated with these four indicators of SES are largely the same.

Next, we used an independent sample of 35,000 participants to address the question of if differences in SES were causal in differences in brain structure and if differences in brain structure were causal in SES differences. We found that total brain volume was one of the many causal factors in differences in occupational prestige, household income, educational attainment, and social deprivation as well as the general factor of SES. We also found that these measures of SES were themselves causal factors in white matter hyperintensities (WMH).

We then looked if differences in cognitive ability accounted for the causal effects of SES differences. We found that for the general factor of SES, once we took account of the effects of cognitive ability, the causal effect of SES on WMH was smaller but still present. This shows that whilst differences in cognitive ability do contribute to the causal effects of differences of SES on WMH other heritable traits also contribute towards the observed effect.

### How can genetic differences be linked to traits like occupation, income, education, and social deprivation?

Whilst at first glance it does seem strange to think that genetic differences are associated with differences in SES. These traits are social variables and so would be influence by things such as the part of the country they live in, the SES of their parents, how the economy is doing, politics and social policies and so on. Undoubtably these factors influence a person’s SES in a myriad of ways. However, in addition to these effects, a person’s SES will be influenced by their own personal characteristics. For example, some of the differences in SES will be linked to differences in health whereby those individuals who are sick less often are better able to attend education and so will likely have a higher SES than those who are sick more frequently. Additionally, those individuals who have a higher level of cognitive ability, or those who are more driven and motivated, will also be more likely to have a higher SES than those who are not. These personal characteristics and traits are heritable meaning that some portion of them is explained by genetic variation. Thus, genetic differences can be linked to differences in SES through other traits that are themselves heritable, a process called vertical pleiotropy (Figure 2.)

**Figure 2.** This figure provides a simplified account of how vertical pleiotropy explains how differences in socioeconomic status can be linked to genetic variation. In instances where differences in socioeconomic status can be explained by personal characteristics such as health, cognitive ability, or personality, then genetic variation linked to differences in these heritable intermediary traits will also be linked with socioeconomic status. Dark blue shows sources of environmental variation with dark blue arrows indicating environmental associations with a trait. Pale blue indicates socioeconomic status. The blue/grey boxes in panel B. show possible intermediary heritable phenotypes.

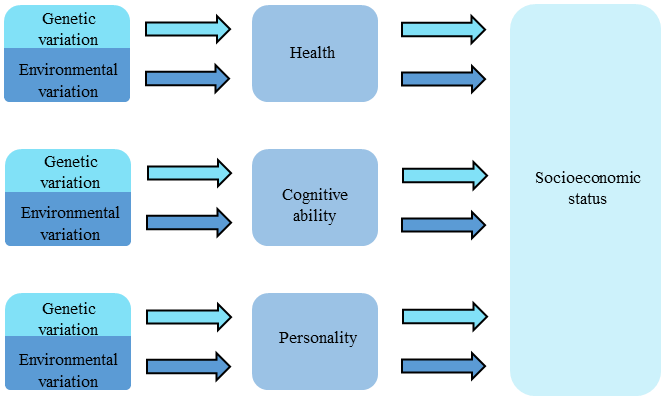

### What is cognitive ability and how does it relate to this study?

As discussed above, our study found evidence that cognitive ability was one of the likely causal factors in differences in SES. We also found evidence that differences in SES were themselves a causal factor in differences in cognitive ability. Indeed, one of the advances made by our study was this bidirectional causal relationship between our measures of SES and cognitive ability. Cognitive ability, often referred to as cognitive function, intelligence or simply as *_g_*, was measured in the current study using a person’s score on a cognitive test, as this is how cognitive ability has been measured in previous large GWAS.

Tests of cognitive ability certainly miss some aspects of human ability that are worthy of study and are not the sole metric by which differences in SES occur. However, for our study we show that scores on this test of cognitive ability are one of the. likely many, causal reasons that individuals differ in their level of SES.

### Did you find the gene for socioeconomic status (occupational prestige, household income, educational attainment, or social deprivation)?

No. Such a characterisation is inaccurate for many reasons. First, SES is a complex trait where many influences act together to produce differences. Some of these potential influences (such as cognitive ability) are themselves heritable. It is possible that some of the genetic associations with factors that influence SES will be picked up using GWAS. Furthermore, there was no gene for SES identified in the sense that there were no large effects found for any individual genetic variant. Rather, many thousands of small genetic effects each make a tiny contribution that in sum acts to influence differences in SES.

### Does this mean genes determine your SES?

No, it does not. In our study we found that a small percentage of differences in SES (and the four indicators of SES) were accounted for by genetic differences. This small association means that even individuals who are similar genetically can end up with a very different SES. However, it does also indicate that there is a slightly higher likelihood that people with a particular genotype will have a higher SES than if they had another genetic combination.

Importantly, the finding that there is a genetic association with a trait does not mean that environmental interventions cannot elicit phenotypic change. One of the classic examples of this is phenylketonuria (PKU). This is a disorder that results in serious medical problems along with intellectual disability. However, despite this disease being genetic in origin, by altering a PKU patients’ diet those afflicted with this condition can lead a life without the symptoms of disease.

The interpretation of the findings of this paper as “genes determine your SES (or any other trait) and there is no way to change this” represents a profound misunderstanding of this study and of genetic research more broadly.

### Could research of this nature lead to discrimination against individuals who have certain genotypes?

Unfortunately, a great deal of scientific research has the potential to be misunderstood and misused. For example, a study aiming to identify genetic variants linked to cardiovascular disease could be abused to discriminate against individuals who have such variants by increasing the cost of their health insurance.

However, it should also be noted that research into the genetic basis of SES can also lead to a decrease in discrimination. This is because the effect size of each individual genetic variant is so small, and the finding that the cumulative effect of all the genetic variants we measured still only accounted for ~11% of the differences in SES.

We also stress that the current study is describing the world as it currently is, not how it could be. Furthermore, our results are specific the countries and cultures that the samples were drawn from and the time in which they were collected. Under different circumstances (i.e. with different sources of environmental variation) the genetic effects that we identify here may become smaller or larger as would the observed differences in peoples SES.

### Can we now tell what someone’s socioeconomic status will be from there genes alone?

No. We found that genetic effects are associated with 11% of occupational prestige, 7% of household income, 13% of educational attainment, 4% of social deprivation and 11% of SES. From these figures we can see that the vast majority of the reasons for differences in these measures of SES is not due to the common genetic variants measured by GWAS.

### What are the practical applications of this research?

At present there are none. We do not advocate any practical application of this work nor draw any policy recommendations from it. This study is a work of basic (as opposed to applied) science, examining links and causes between variables of interest. The study did not include an intervention or any attempt to improve a trait.
